# Presenting the *Compendium Isotoporum Medii Aevi* (CIMA) and Bayesian Case Studies

**DOI:** 10.1101/2021.08.05.455253

**Authors:** Carlo Cocozza, Enrico Cirelli, Marcus Groß, Wolf-Rüdiger Teegen, Ricardo Fernandes

## Abstract

The *Compendium Isotoporum Medii Aevi* (CIMA) gathers more than 50 000 isotopic measurements for bioarchaeological samples located within Europe and its margins dating between AD 500–1500. This volume of isotopic data, together with collected supporting information, offers multiple research opportunities. This is illustrated here using novel Bayesian modelling methods on selected case studies to reconstruct medieval human lifeways (i.e. human subsistence, spatial mobility), animal management practices, and paleo-environmental conditions. We also discuss how the integration of isotopic data with other types of archaeological and historical data can improve our knowledge of historical developments throughout medieval Europe.

## Introduction

In the late 1970’s, stable carbon isotope analysis was first employed for paleo-dietary reconstruction through the analysis of human remains (Vogel & Van der Merwe 1977). Since then, the use of isotopic methods in archaeological research has expanded following several developments in isotope ratio mass spectrometry methods and lab pre-treatment protocols that increased the number of measurable isotopic ratios across a wider variety of materials (Leng 2006; Meier-Augenstein 2011; Richards & Britton 2020). Such developments have allowed for a larger number of applications in archaeological research and for more accurate and precise assessments of past phenomena. The reconstruction of past human subsistence, nutrition and spatial mobility, the study of past animal and crop management practices, or the reconstruction of paleo-environments and -climates are just some examples that illustrate the importance of isotopic methods in archaeological research (Hedges *et al.* 2004; Lee-Thorp 2008; Fiorentino *et al.* 2015; Lightfoot & O’Connell 2016). This is also evident from the exponential growth in recent decades in the number of archaeological publications reporting isotopic results (Roberts *et al.* 2018). Once collected and curated, amassed isotopic data can be subject to meta-analyses from which it is possible to investigate past human and natural phenomena at varying spatial and temporal scales (Cubas *et al.* 2020; Wilkin *et al.* 2020; Wang *et al.* 2021).

The Middle Ages (c. fifth to fifteenth centuries AD) is a formative period of European history. It was marked by major transformations in political and economic systems, vast population movements, violent armed conflicts, climate change, development of religious movements, and technological innovations, albeit with regional variations (Brown 1971; Holmes 1988; Backman 2003; Wickham 2016). The study of such historical phenomena has been predominantly based on written sources although these may vary in quality and representation. In particular, the lifestyles of lower socioeconomic classes are often mis- or under-represented given their illiteracy (Baten & Steckel 2019). Such knowledge gaps can be reduced by isotopic analyses of human remains from which it becomes possible to build iso-biographies describing the diets and spatial mobility of single individuals from across socioeconomic, religious, and cultural spectra (e.g. Lamb *et al.* 2014; Alexander *et al.* 2015; Hughes *et al.* 2018). Isotopic analyses of animal and plant remains have also been employed in medieval contexts to reconstruct past climatic and environmental conditions and economic and agricultural activities (e.g. Hamilton & Thomas 2012; Reitsema *et al.* 2013; Hamerow *et al.* 2020). To explore the research opportunities offered by published isotopic data concerning the European Middle Ages we compiled the open-access CIMA (*Compendium Isotoporum Medii Aevi*) database. The database is part of the Pandora & IsoMemo initiatives that bring together networks of autonomous databases involved in the study of the human past in addition to tools for data access and advanced modelling. Here we describe CIMA and illustrate its research potential through the application of Pandora & IsoMemo modelling in different case studies.

### Database structure and data selection criteria

Data collected for CIMA dates from the fifth to the late fifteenth centuries. Data spatial coverage includes all of Europe, but also non-European regions presenting cultural or religious affinities (e.g. Norse populations in Greenland or Christian Crusaders in Jordan or Palestine). Our compilation includes isotopic measurements on human, animal, and plant remains for different tissue types plus other analytical data to support their interpretation (e.g. atomic C/N ratios). We also included supporting information describing the archaeological sites from which samples originated and their location and chronology. In the case of human data, whenever available we included osteological data (e.g. estimated age) and likely socioeconomic status and religion. A description of the database structure is given in Supplementary Information file S1.

### Description of collected data

At time of writing (May 2021) the database consists of 17 756 human, 4946 animal and 164 plant entries. This data was collected from 358 primary sources which included journal articles, book chapters, archaeological reports, and academic dissertations. The total number of carbon, nitrogen, sulphur, strontium, and oxygen isotopic measurements included in the database is 50 153. Most of the collected data is from archaeological sites located in the UK (24.1%), followed by Italy (10.8%), Spain (9.6%), and Germany (8.0%). The spatial distribution of archaeological sites for data sources is shown in Fig. 1. It shows a major data gap for France (3.1% of data) which is compounded by its size and importance in medieval European history. Additional summaries and descriptions of human, animal, and plant data can be found in Supplementary Information file S1.

**Fig.1.**
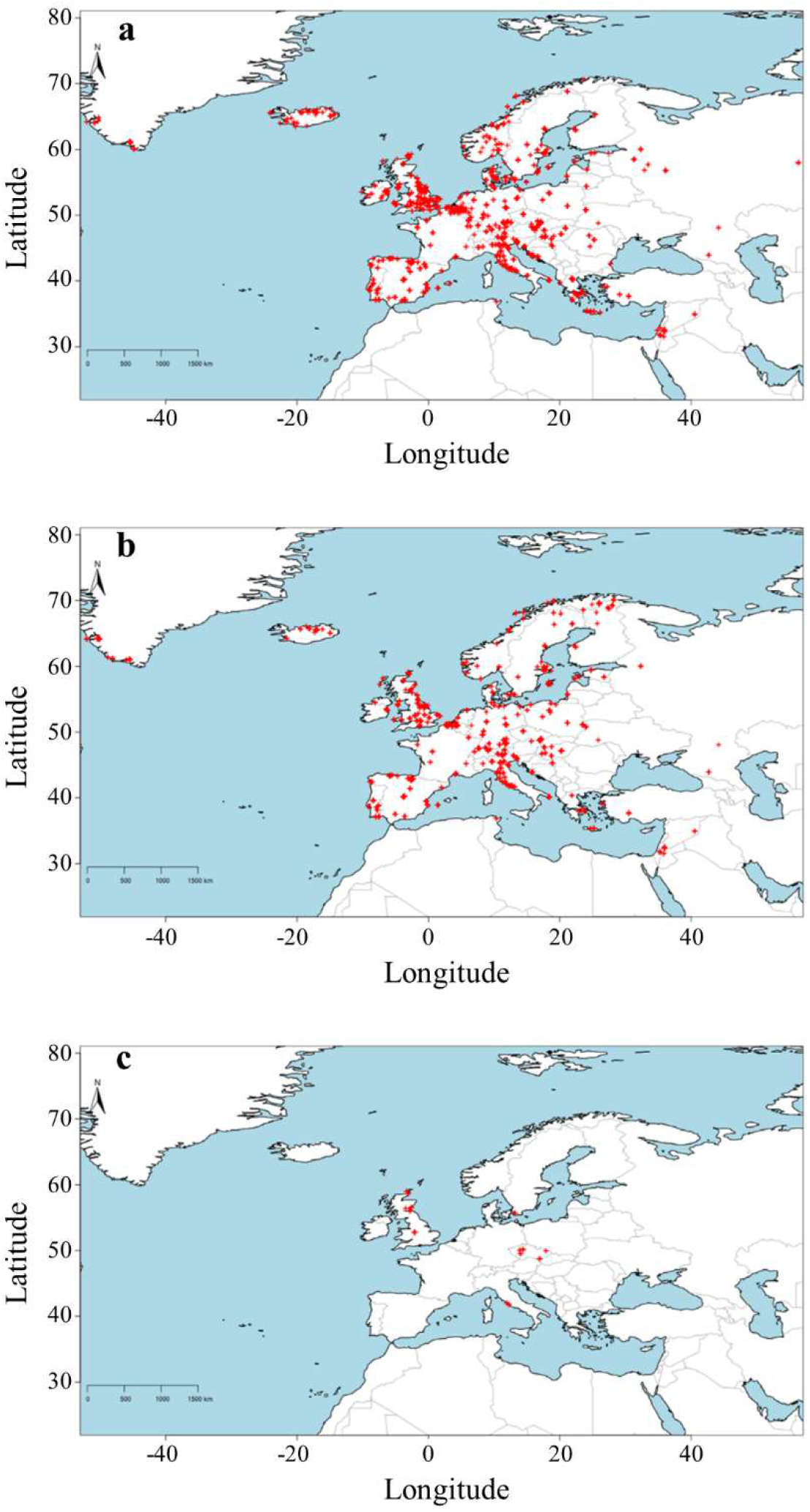
Spatial distribution of human (a), animal (b), and plant (c) site locations for data compiled within CIMA.

CIMA is a partner database of and was developed within the Pandora & IsoMemo initiatives that established networks of independent databases for the study of the human past and isotopes across time, respectively. The database (https://doi.org/10.48493/s9nf-1q80) is made available via MATILDA: a repository for Medieval bioAnThropologIcaL DatAbases (https://pandoradata.earth/organization/matilda-a-repository-for-medieval-bioanthropological-databases). MATILDA is itself integrated within the Pandora and IsoMemo Big Data initiatives. CIMA is designed as a collaborative effort and we welcome future data contributions from other researchers. We have implemented a system through which separate dataset contributions are assigned a unique DOI. These datasets are then included in a master file that combines all data and includes fields to reference primary data sources plus data compilations. This provides a chain of data and ensures that efforts for both data production and data collection are recognised.

### Bayesian modelling

The Pandora and IsoMemo initiatives are also engaged in the development of R-based modelling tools which are made available online via a user-friendly interface (https://www.pandoraapp.earth). This includes, among others, Bayesian spatio-temporal modelling of isotopic data, the tracking of human or animal spatial mobility, and dietary reconstruction (Fernandes *et al.* 2014; Cubas *et al.* 2020; Wilkin *et al.* 2020; Cocozza *et al*. 2021; Wang *et al.* 2021). We present here illustrative case studies to present the research potential offered by Pandora & IsoMemo modelling capabilities on CIMA data. Detailed descriptions of employed modelling methods are given in Supplementary Information file S2.

*Case study 1: Distribution of animal isotopic values in medieval Europe: animal management, paleo-climatic and -environmental conditions, and baselines for dietary studies*

Types of local vegetation cover and plant δ^13^C and δ^15^N values are influenced by several factors such as water availability, canopy effects, altitude, soil δ^15^N, to mention only a few, in addition to human crop management practices such as irrigation or manuring (Styring *et al.* 2016). The isotopic values of domesticated herbivores reflect those of consumed vegetation while domesticated omnivores may also consume animal protein. In addition, domesticated animals may be subject to selective feeding and livestock fencing. Thus, isotopic values from archaeological animal remains are a palimpsest of information on animal management practices and local environmental/climatic conditions. To illustrate this research potential, we here make a broad diachronic (Iberia, Italy, and England) and spatial (Europe) comparison of isotopic values for domesticated herbivores (cattle/ovicaprids) and omnivores (pigs/chickens).

The diachronic comparison (time bin AD 500 to 1000 vs. AD 1000 to 1500) for selected regions (Fig. 2 and 3) and the observed spatial pattern for all combined periods (Fig. 4) show that Italy and Iberia have roughly similar patterns for both δ^13^C and δ^15^N and that these differ from England when it comes to domesticated herbivores. In the case of herbivore δ^13^C, to an extent this reflects a higher water abundance and greater canopy effect in northern Europe but some of the more elevated δ^13^C values in southern Europe reveal some animal consumption of C_4_ plants (e.g. millet, sorghum) or in the case of Muslim Iberia, also sugarcane productions wastes (Montanari 1988; Castiglioni & Rottoli 2013; Alexander *et al.* 2015). The δ^15^N values for herbivores in England show narrower ranges than those in Italy/Iberia but are similar for omnivores, in spite of the considerably larger environmental variability in Iberia/Italy. Given that there are no significant temporal differences, this suggests that omnivores feeding and crop/vegetation management practices differed across territories within medieval England (Rippon *et al.* 2014).

**Fig.2.**
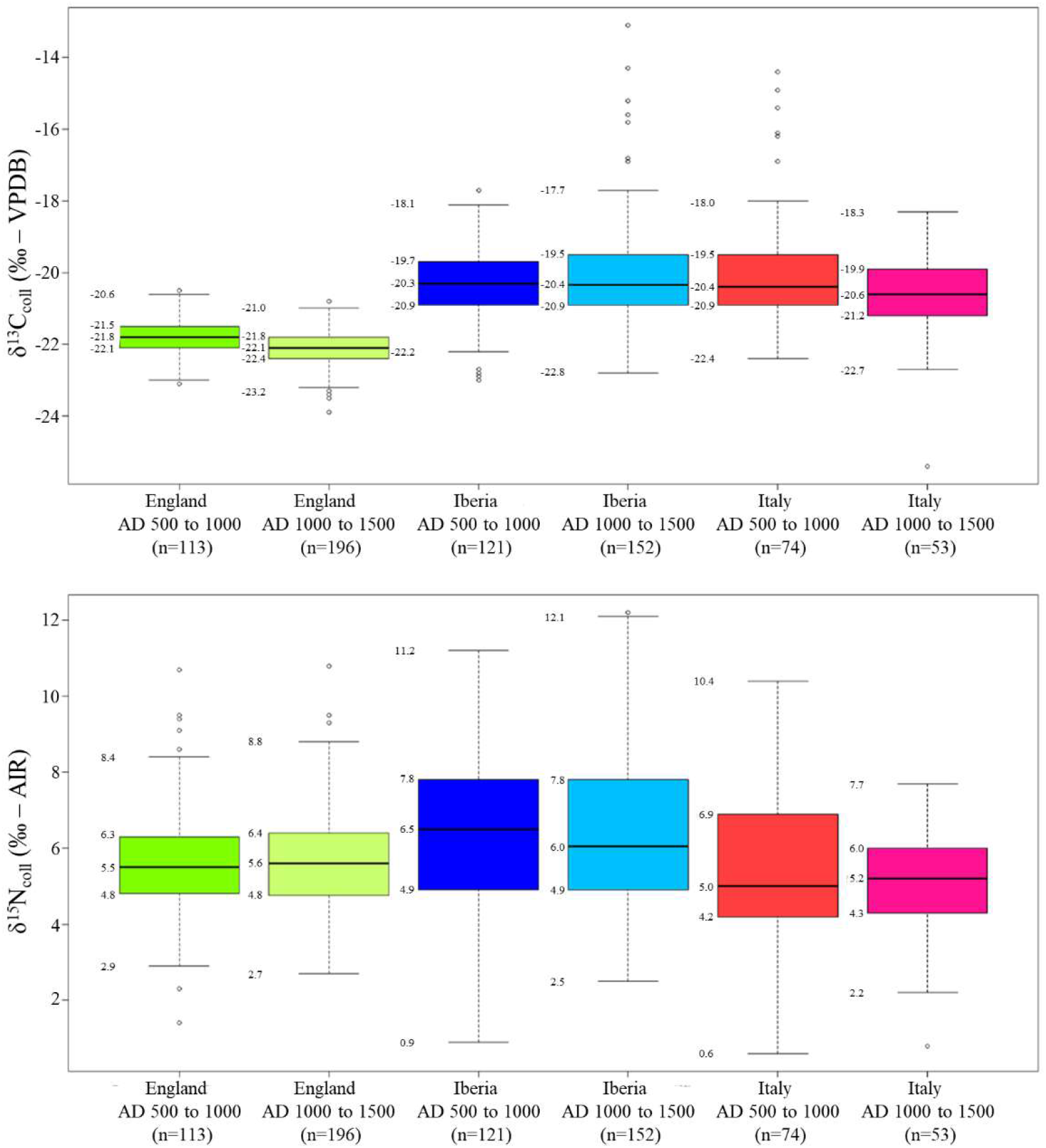
Spatio-temporal comparison of δ^13^C and δ^15^N bone collagen values from domesticated herbivores. Box edges represent the 25^th^ and 75^th^ percentiles (inter quartile range or IQR) while the horizontal line is the 50^th^ percentile (median). Whiskers are the minimum and maximum values that do not exceed a distance of 1.5 times the IQR. Beyond this distance single points represent outliers.

**Fig.3.**
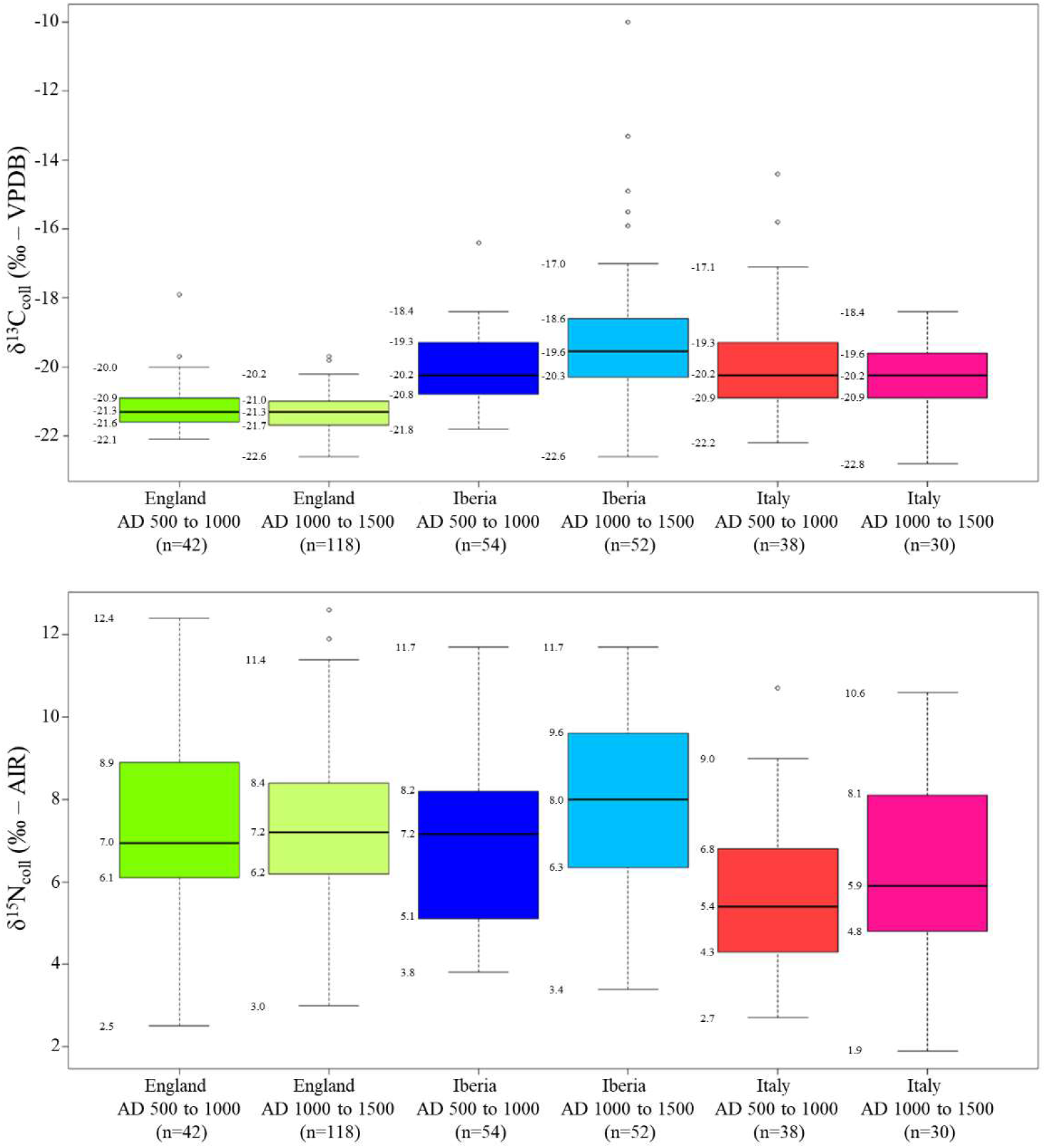
Spatio-temporal comparison of δ^13^C and δ^15^N bone collagen values from domesticated omnivores. Box edges represent the 25^th^ and 75^th^ percentiles (inter quartile range or IQR) while the horizontal line is the 50^th^ percentile (median). Whiskers are the minimum and maximum values that do not exceed a distance of 1.5 times the IQR. Beyond this distance single points represent outliers

**Fig.4.**
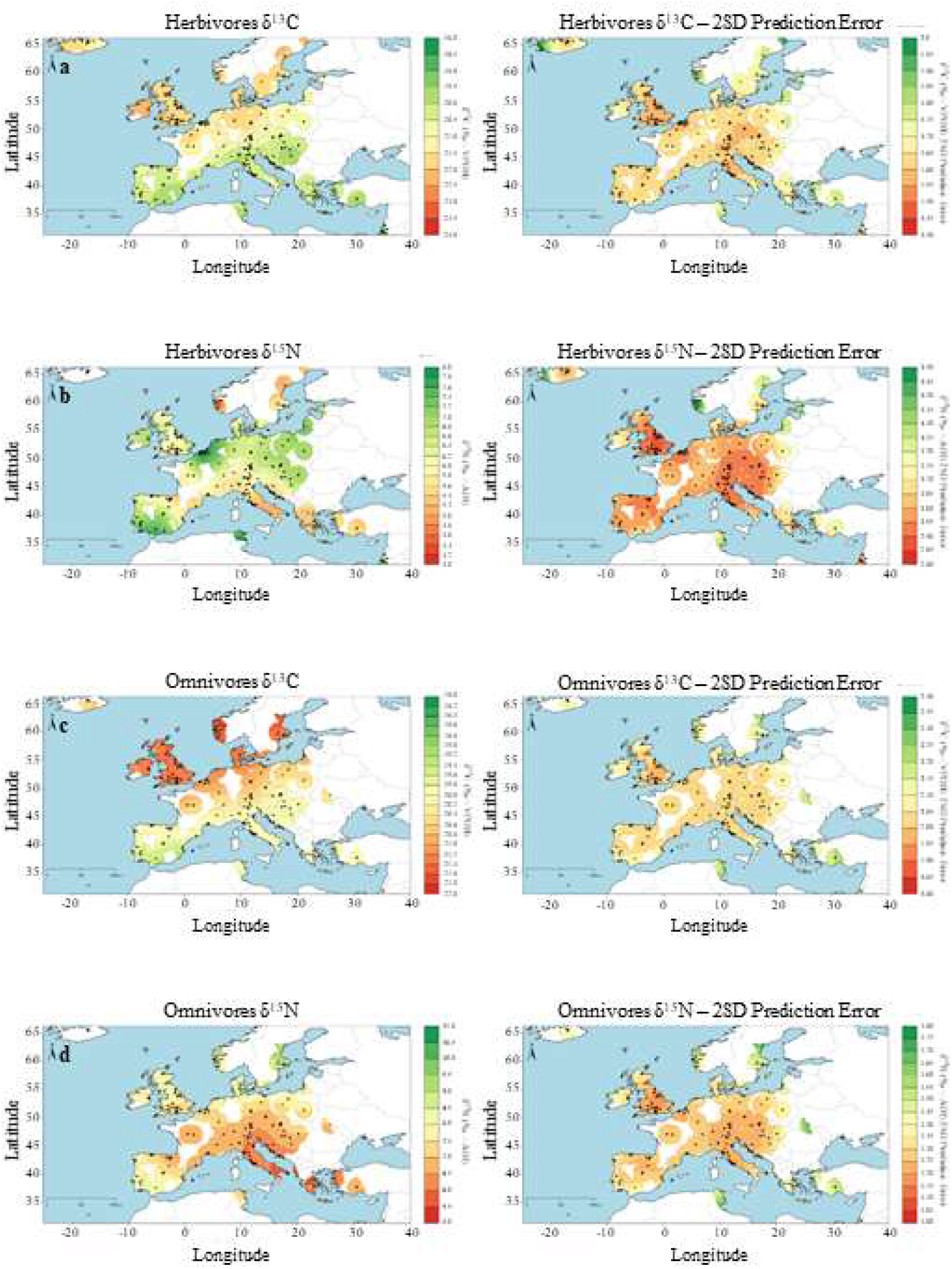
Spatial comparison of δ^13^C and δ^15^N mean and prediction errors for domesticated animals. a) δ^13^C herbivores; b) δ^15^N herbivores; c) δ^13^C omnivores; d) δ^15^N omnivores.

*Case study 2: Human diets during the first millennium AD: spatial and temporal isotopic trends, Bayesian dietary reconstruction, and Bayesian integration of isotopic, historical, archaeobotanical, and archaeofauna data*

The collapse of the Western Roman Empire (AD 476), the splitting of its territories into separate kingdoms, and the subsequent territorial unification attempt during the Carolingian empire (AD 800– 887) mark major historical transitions in Europe. Different sources of historical and archaeological evidence point towards a high diversification of farming and animal rearing in Late Roman to early medieval Europe, yet far from the intensive agricultural economy of the Roman empire (Montanari 1988; Lewit 2009; Witcher 2016; MacKinnon 2019). In concomitance, the arrival of migrating populations may have also shifted dietary habits (Pearson 1997). We combined CIMA and IsoArcH isotopic data, a database for the Greco-Roman world (Salesse *et al.* 2018), to map and compare spatial distribution of human adult bone collagen carbon (δ^13^C) and nitrogen (δ^15^N) stable isotopes for three time slices: AD 200, AD 500, and AD 800. From this we excluded individuals identified as high-status to avoid a potential bias related to preferential consumption of high value foods. Variations in human δ^13^C and δ^15^N may reflect differences in the proportions of consumed protein sources with distinct isotopic values and/or shifts in the isotopic values of consumed foods resulting from changes in climatic/environmental conditions and/or in crop or animal management practices (Hedges *et al.* 2004; Lee-Thorp 2008; Fiorentino *et al.* 2015).

Figures 5–6 show the spatial average difference for adult bone collagen δ^13^C and δ^15^N for adjacent time slices and whether these are statistically significant (p<0.1) (Fig. 5–6; also S3-S4, Supplementary Information). There is significant increase in δ^13^C for Galicia (northern Spain), Northern Italy and in the former Roman province of Pannonia (northern Balkans) but these do not show significant differences for δ^15^N. This suggests an increase in the consumption of C_4_ plants (e.g. millet and/or sorghum) and/or products from animals foddered on these. The break in the Roman economic and agrarian system reduced access to wheat and barley, whereas millet and sorghum became commonly consumed by the lower classes in northern Italy and in the Balkans (Montanari 1988; Winklerová 2011; Castiglioni & Rottoli 2013; Gyulai 2014). In Galicia, the arrival of the Germanic Suebi and environmental shifts possibly led to an increase in C_4_ consumption (López-Costas & Müldner 2016). Significant decreases in δ^15^N are observed in northern France and southern England which could result from a lower consumption of animal protein sources and/or a shift in food isotopic values arising from changes in agricultural practices (e.g. lower cereal manuring intensity) (Lewit 2009).

**Fig.5.**
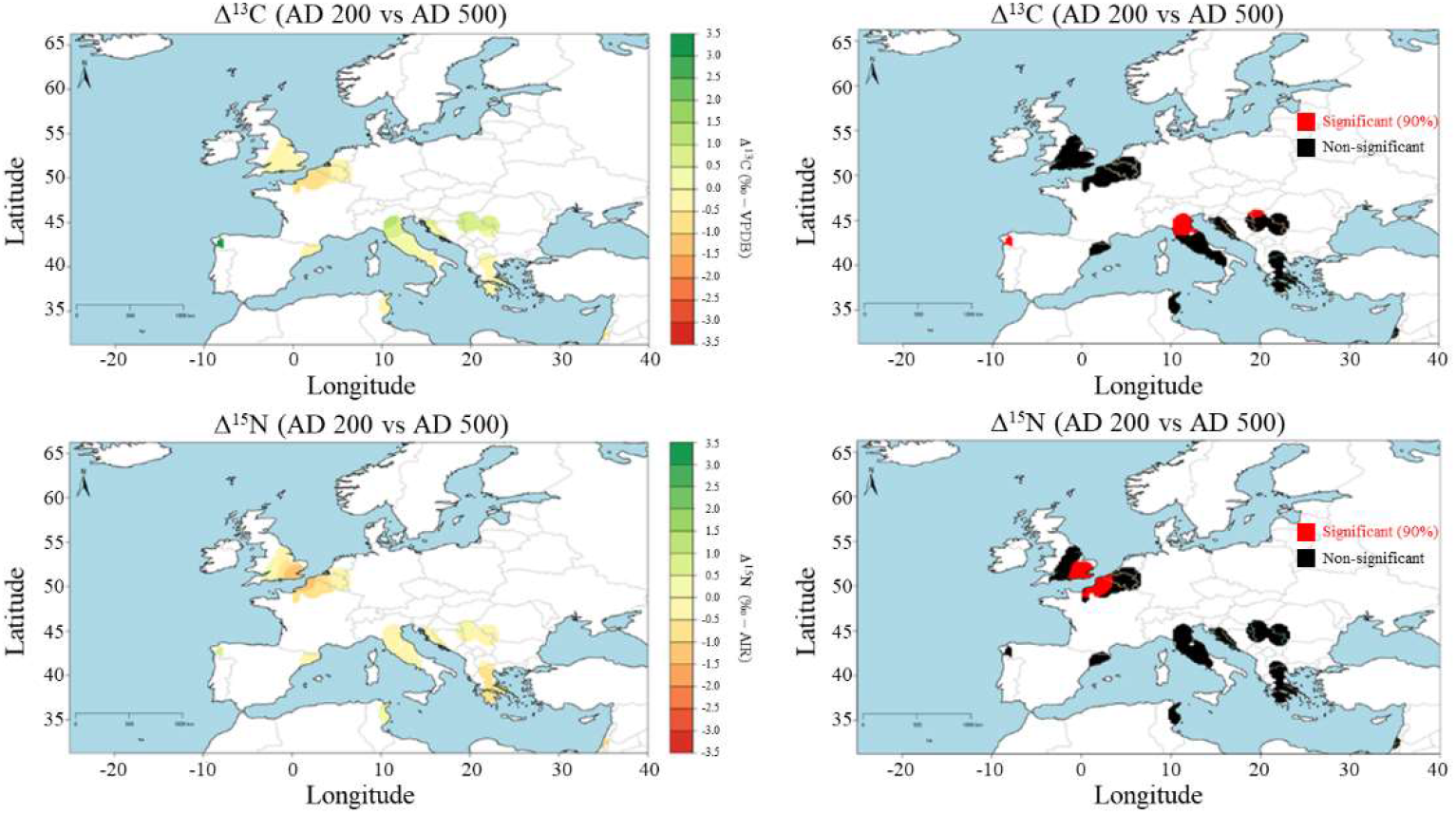
Left column: Bayesian mapping of differences in average human bone collagen isotopic values (Δ^13^C and Δ^15^N) for two time slices (AD 200 vs. AD 500). Right column: statistical significance of difference (p<0.1).

**Fig.6.**
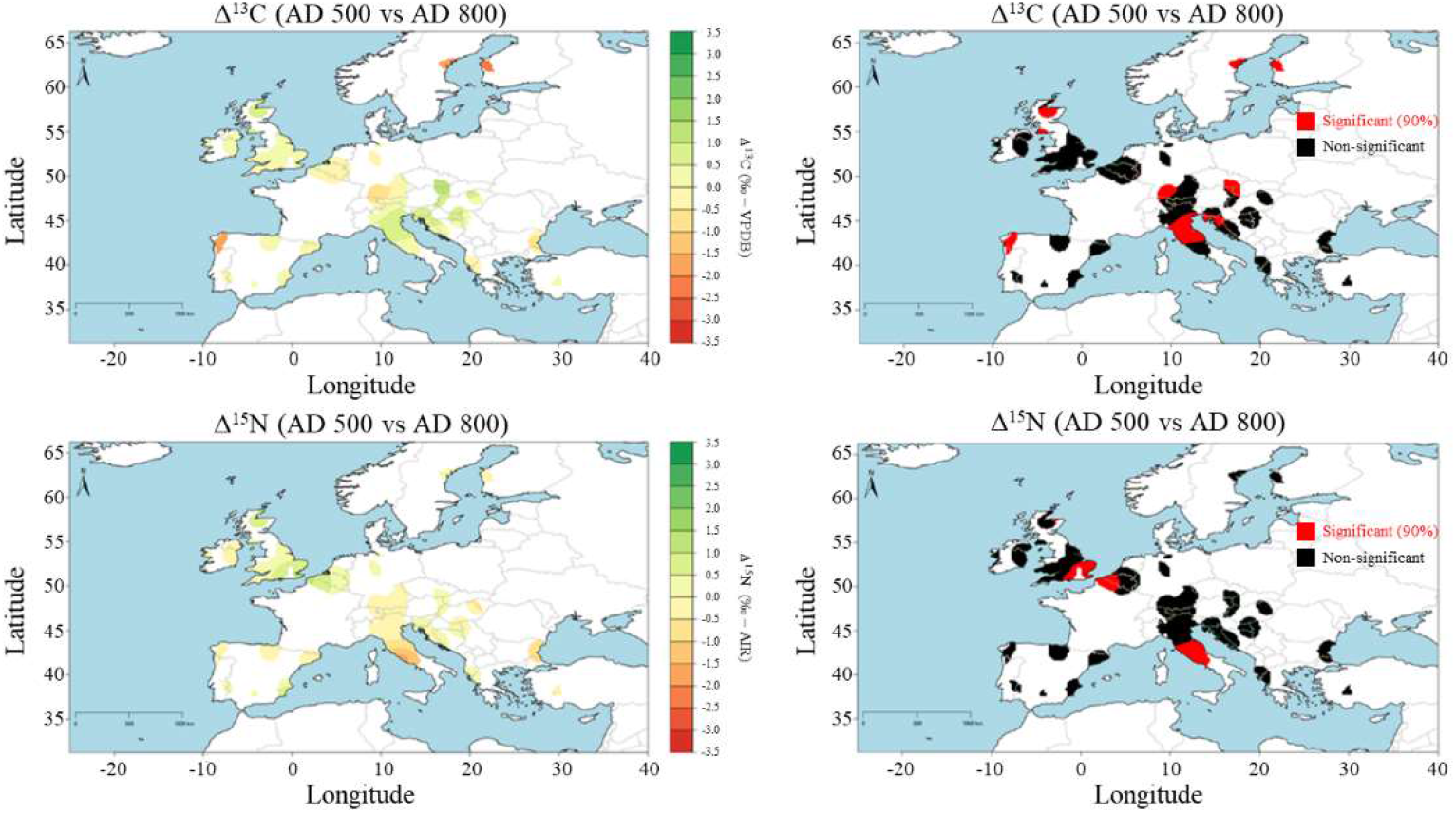
Left column: Bayesian mapping of differences in average human bone collagen isotopic values (Δ^13^C and Δ^15^N) for two time slices (AD 500 vs. AD 800). Right column: statistical significance of difference (p<0.1).

The comparison of the AD 500 and AD 800 time slices reveals regions with significant shifts, in some cases overlapping with regions discussed above. In the case of northern Italy and the Balkans, the continued increase in δ^13^C values suggests that the cultivation of C_4_ cereals gradually grew during the early medieval period (Montanari 1988; Winklerová 2011; Castiglioni & Rottoli 2013; Gyulai 2014). In central Italy, there is a decrease in δ^15^N average values. Here a reduction in animal sizes (Salvadori 2019) and a general shift towards silvopastoralism (MacKinnon 2019; Salvadori 2019) would be consistent with a decline in the consumption of terrestrial animal protein and/or a decrease in animal δ^15^N values as consequence of free-roaming rearing practices.

To showcase Bayesian modelling of the temporal variability in human bone collagen isotopic values at one location we selected the city of Rome between AD 1 and 1000 (Fig 7). The model produced for both sexes shows relatively constant δ^13^C values and an overall decrease in δ^15^N values for adults albeit with some variability. Isotopic ranges for both sexes greatly overlap and do not reveal statistically significant differences. Some historical sources suggest the existence of gender-based nutritional inequality in antiquity, although their extension beyond restricted communities (e.g. monastical) is unknown (Bynum 1987; Pearson 1997; Garnsey 1999). The overall decrease in human δ^15^N values likely reflects a combination of factors, including the end of the Roman proto-welfare system (*Annona* i.e. the yearly distribution of grain in Rome, which at times included pork) and a reduction in the proportion of consumed pork in favour of ovicaprids as revealed by archaeofaunal studies.

**Fig.7.**
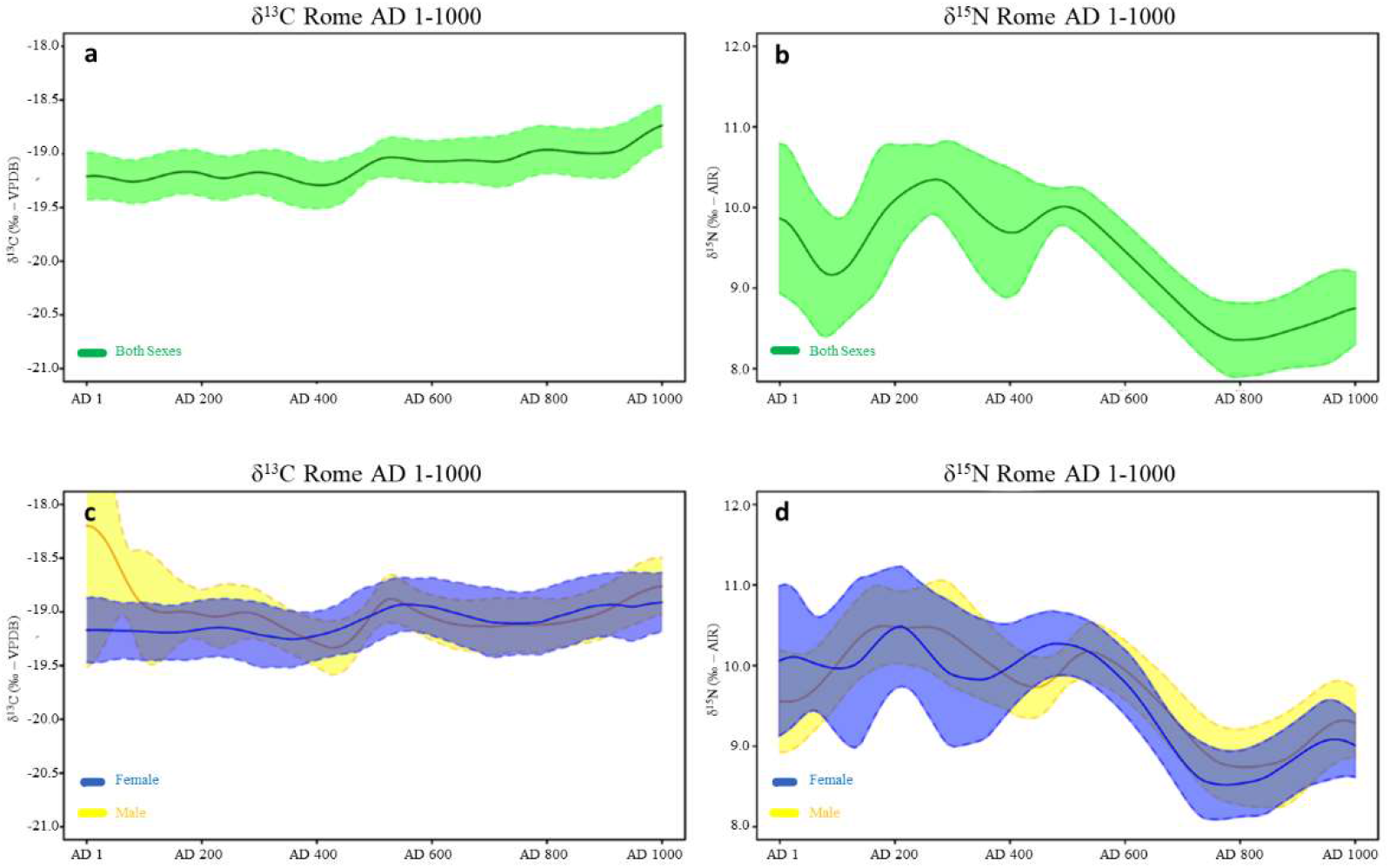
Temporal Bayesian plots for adult bone collagen δ^13^C and δ^15^N values for Rome. a) both sexes δ^13^C; b) both sexes δ^15^N; c) female *versus* male δ^13^C; d) female *versus* male δ^15^N.

To obtain dietary estimates for lower status adults in Rome during Roman Imperial and medieval periods we employed the Bayesian software ReSources, an upgraded version of the FRUITS (Fernandes *et al.* 2014) (modelling details in Supplementary Information file S2). Modelled data consists of human and food isotopic values and as shown in Fig. 8 (left) it was not possible to obtain precise dietary estimates given the uncertainties in model parameters and issues of equifinality. That is, when varying proportions of food contributions generate the same human isotopic value. To overcome this, we explored the use of prior dietary information from written, archaeobotanical, and archaeofaunal studies to define a dietary scenario matching the assemblage of data. Such studies often offer semi-quantitative relationships that can be used under a Bayesian approach to set conservative boundaries on food contributions or express inequality relationships among food contributions, both synchronically and diachronically (Fernandes *et al.* 2015). The defined dietary scenario allowed us to increase the precision of dietary estimates for caloric contributions, macronutrient intakes, and protein contributions which can be compared with modern population results (Fig. 8, right and Supplementary Information file S5). In all three time-slices the consumption of wheat products is predominant among all the main food sources, although this decreases through time. Among animals, pigs are the main source of calories. The major difference between modern and ancient populations is that the latter consumed higher amounts of pulses.

**Fig.8.**
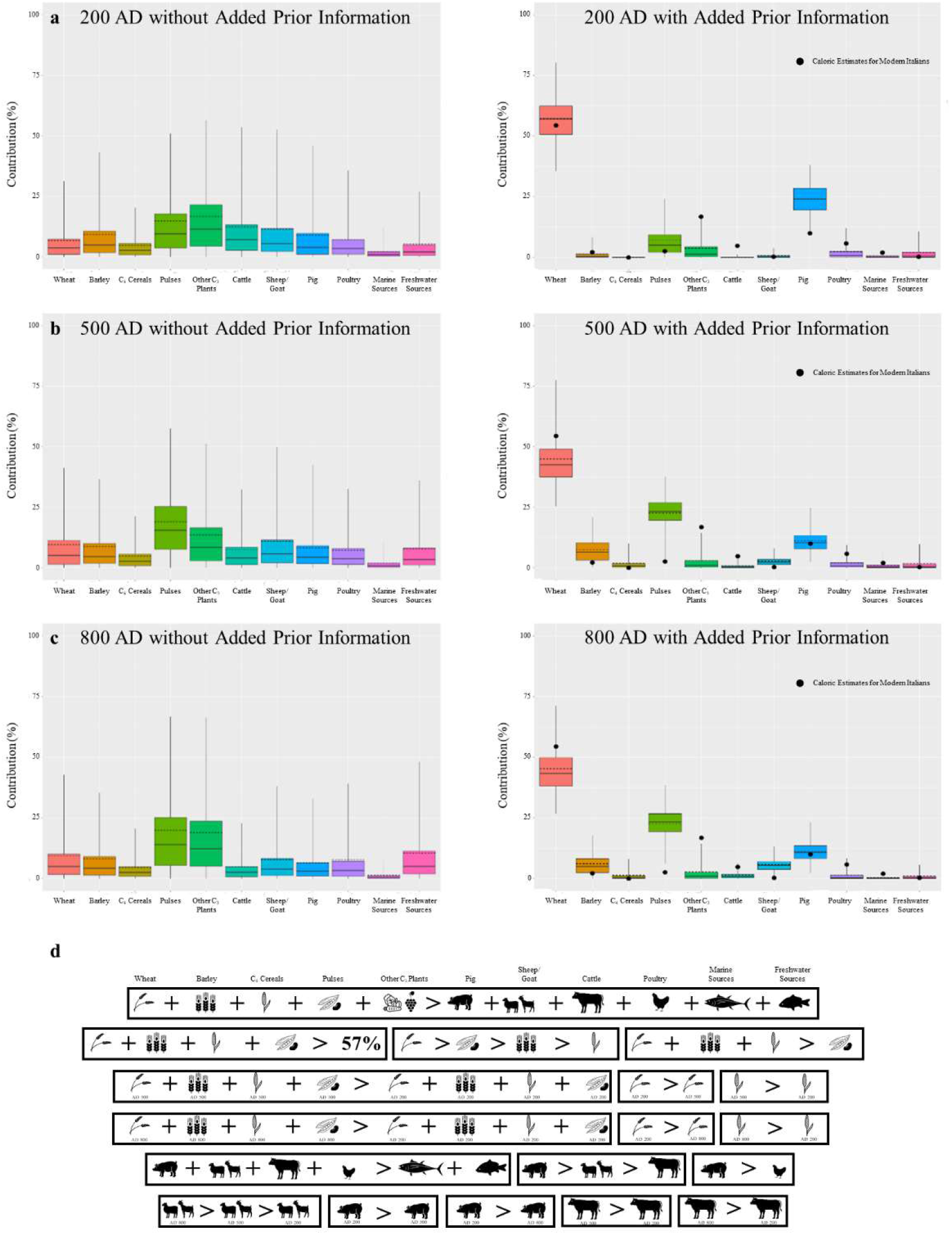
AD 200 (a), 500 (b) and 800 (c) dietary estimated models of main food sources caloric contribution for the city of Rome without (left) and with (right) added prior dietary information (d). Black circles within plots correspond to modern dietary estimates. See also Supplementary Information files S2 and S5.

#### Example 3: Human spatial mobility in Roman and medieval Britain

Medieval Europe witnessed several population movements at various scales, from the mass migrations of the Germanic Migration Period (conventional AD 375–568), to comparatively smaller scale movements following military conflicts, urbanisation processes, and religious pilgrimages (Backman 2003; Wickham 2016). Isotopic studies of human mobility typically involve the use of isotopic ratios of strontium (^87^Sr/^86^Sr) or oxygen (δ^18^O) which vary spatially in available water. These isotopic signals are measured in human tissues with varying formation periods and turnover rates. and, from comparisons with isotopic ranges in the vicinity of burial location, an individual may be categorised as mobile or non-mobile. In cases where spatial variations in isotopic values are known for wider regions it may be possible to assign a place of residence for mobile individuals.

Here we investigated mobility patterns for Roman and medieval individuals buried at sites in York and London. We measured ^87^Sr/^86^Sr or δ^18^O on teeth with a Bayesian baseline for each isotopic proxy (details in Supplementary Information file S2). Individuals for which the values for one of these proxies did not match local values were categorized as mobile or otherwise as non-mobile. In assessing temporal trends in spatial mobility, we compared kernel density plots for total population subject to either ^87^Sr/^86^Sr or δ^18^O analysis, mobile individuals, and non-mobile individuals. Relative variations in kernel density values for these can be used to assess variations in the proportion of mobile populations at burial sites. However, it is important to notice that this study is subject to a potential sampling bias given that isotopic mobility studies often target preferentially cemeteries that are expected mobile individuals. The comparison with more frequent dietary isotopic analyses (Fig. 9a) shows that mobility studies are more concentrated in specific time periods (Roman and late medieval) although data gaps are also present in dietary studies. These may result not only from sampling bias but also from sample availability which depends on past population numbers, burial practices, and taphonomic effects.

**Fig. 9.**
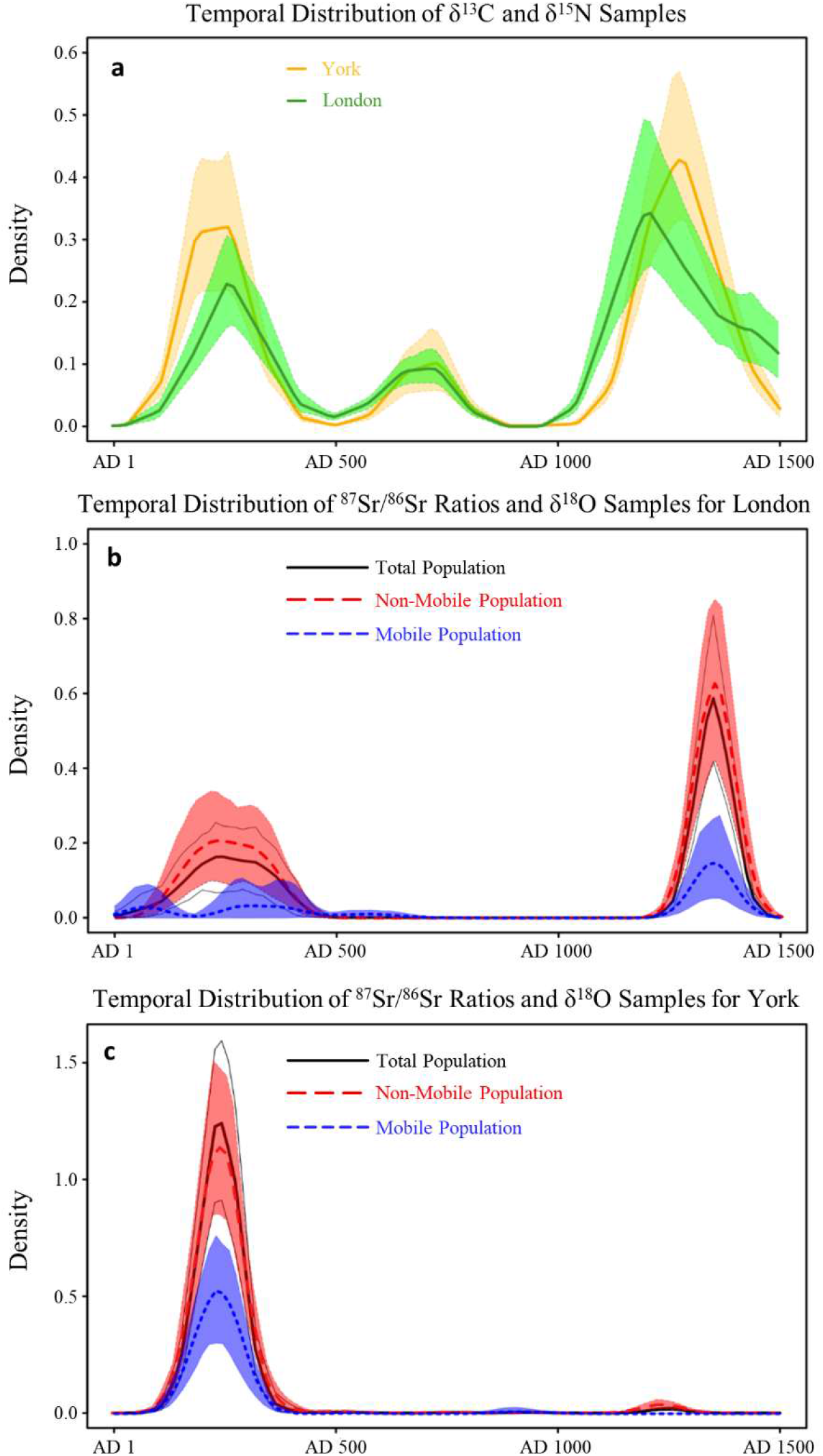
Kernel density plots for human osteological samples from London and York.

Kernel density plots comparing mobile and non-mobile individuals reveal some changes in their proportions relative to the total analysed population (Fig. 9). The kernel density results for London, show that the proportion of mobile individuals is comparatively higher during the early Roman Period and during the Anglo-Saxon migration in the fifth century AD. There are also mobile individuals dating to the late medieval period although in a comparatively smaller proportion, since these correspond to a study on the Black Death. The Roman population of York also contained several mobile individuals and a small peak shows one mobile individual dating to the ninth century of probable Scandinavian origins.

The place of origin of a mobile individual can be estimated by comparing ^87^Sr/^86^Sr and/or δ^18^O values measured in a tooth formed at an early age (under the assumption that the individual spent early age at a certain location) with spatial maps for these isotopic proxies. The recently developed LocateR model can then be used to generate probability density maps on the place of origin of the individual (Wang *et al.* 2021). Relying on reported δ^18^O values we employ this approach to estimate the place of origin for three individuals (REP-295, REP-511, REP-529) buried in Repton, UK and associated with the Scandinavian ‘Great Heathen Army’, invading Britain in the late ninth century (Biddle & Kjølbye-Biddle 2001; Budd *et al.* 2004; Jarman *et al.* 2018). We also estimated, using a combination of^87^Sr/^86^Sr and/or δ^18^O measurements, the place of origin of a young individual (SK27), buried in a high medieval leprosarium in Winchester, UK with a scallop shell typical of a pilgrim who completed the travel to Santiago de Compostela. This individual was previously found to be non-local but likely still native to Britain (Roffey *et al.* 2017).

Results (Fig. 10) show that the two Repton individuals buried in a double grave (REP-295 and REP-511) likely originate from the British Isles or the continental coasts from Normandy to south-western Denmark. Interestingly, the highest probability would point towards Ireland, which is often mentioned in historical sources as associated with campaigns led by some of the leaders of the army. The other Repton individual (REP-529) has a likely place of origin in Sweden, Norway, and the Baltic region, in accordance with the archaeological evidence. Results for the pilgrim buried in Winchester suggest that he was likely residing in northern England or in southern Scotland during tooth formation.

**Fig. 10.**
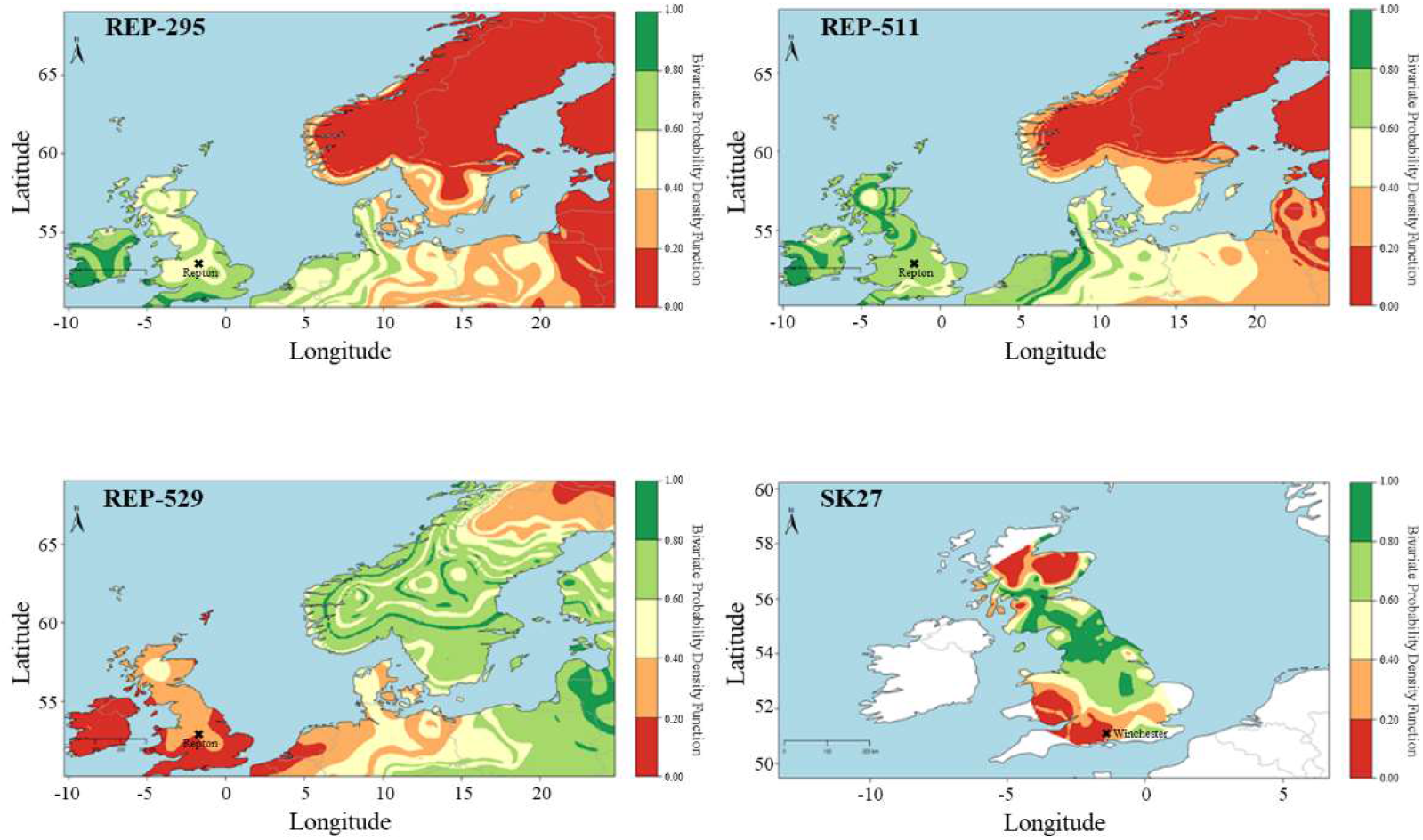
Probability density maps for place of origin for individuals REP-295, REP-511, REP-529 and SK27.

## Conclusion

This paper presented the *Compendium Isotoporum Medii Aevi* (CIMA), the largest archaeological isotopic database on medieval Europe. This dataset can be employed for meta-analyses studies of human lifeways, agricultural practices, and paleo-environmental/climatic conditions. It can also support future isotopic studies by providing reference values or used to identify data gaps requiring additional research. CIMA data compilation efforts will continue and the initiative is open to collaborations with other researchers.

## Acknowledgements

The data was collected as part of the Pandora & IsoMemo initiatives supported by Max Planck Institute for the Science of Human History, PS&H research group, University of Warsaw, Masaryk University, and Eurasia3angle research group. We thank all researchers who have published stable isotope measurements on Medieval Europe and contributed indirectly to the formation of this collection.

## References

Alexander, M.M., C.M. Gerrard, A. Gutiérrez & A.R. Millard. 2015. Diet, Society, and Economy in Late Medieval Spain: Stable Isotope Evidence From Muslims and Christians from Gandía, Valencia. American Journal of Physical Anthropology, 156: 263–273. https://doi.org/10.1002/ajpa.22647

Backman, C.R. 2003. The Worlds of Medieval Europe. Oxford: Oxford University Press.

Baten, J. & R.H. Steckel. 2019. The History of Violence in Europe, in R.H. Steckel, C.S. Larsen, C.A. Roberts & J. Baten (ed.) Backbone of Europe. Health, Diet, Work and Violence over Two Millennia: 300–324. Cambridge: Cambridge University Press.

Biddle, M. & K. Kjølbye-biddle. 2001. Repton and the ‘great heathen army’, 873–4, in J. Graham-Campbell, R. Hall, J. Jesch & D.N. Parsons (ed.) Vikings and the Danelaw: 45–96. Oxford: Oxbow Books.

Brown, P. 1971. The World of Late Antiquity. London: Thames & Hudson.

Budd, P., A. Millard, C. Chenery, S. Lucy & C. Roberts. 2004. Investigating population movement by stable isotope analysis: a report from Britain. Antiquity, 78: 127–141. https://doi.org/10.1017/S0003598X0009298X

Bynum, C.W. 2010. Holy Feast and Holy Fast. The Religious Significance of Food to Medieval Women. Berkeley: University of California Press.

Castiglioni, E. & M. Rottoli. 2013. Broomcorn millet, foxtail millet and sorghum in north Italian Early Medieval sites. European Journal of Post Classical Archaeologies, 3: 131–144.

Cocozza, C., R. Fernandes, A. Ughi, M. Groß & M.M. Alexander. 2021. Investigating infant feeding strategies at Roman Bainesse through Bayesian modelling of incremental dentine isotopic data. International Journal of Osteoarchaeology. https://doi.org/10.1002/oa.2962

Cubas, M. ET AL. 2020. Latitudinal gradient in dairy production with the introduction of farming in Atlantic Europe. Nature communications, 11: 2036. https://doi.org/10.1038/s41467-020-15907-4

Fernandes, R., A.R. Millard, M. Brabec, M.J. Nadeau & P. Grootes. 2014. Food Reconstruction Using Isotopic Transferred Signals (FRUITS): A Bayesian Model for Diet Reconstruction. PLoS ONE, 9: e87436. https://doi.org/10.1371/journal.pone.0087436

Fernandes, R, P. Grootes, M.J. Nadeau & O. Nehlich. 2015 Quantitative diet reconstruction of a Neolithic population using a Bayesian mixing model (FRUITS): the case study of Ostorf (Germany). American Journal of Physical Anthropology, 158: 325–340. https://doi.org/10.1002/ajpa.22788

Fiorentino, G., J.P. Ferrio, A. Bogaard, J.L. Araus & S. Riehl. 2015. Stable isotopes in archaeobotanical research. Vegetation History and Archaeobotany, 24: 215–227. https://doi.org/10.1007/s00334-014-0492-9

Garnsey, P. 1999. Food and Society in Classical Antiquity. Cambridge: Cambridge University Press.

Gyulai, F. 2014. The history of broomcorn millet *(Panicum miliaceum* L.) in the Carpathian-basin in the mirror of archaeobotanical remains II. From the Roman age until the late medieval age. Columella: Journal of Agricultural and Environmental Sciences, 1. DOI: 10.18380/SZIE.COLUM.2014.1.1.39

Hamerow, H., A. Boogard, M. Charles, E. Forster, M. Holmes, M. Mckerracher, S. Neil, C. Bronk Ramsey, E. Stroud & R. Thomas. 2020. An Integrated Bioarchaeological Approach to the Medieval ‘Agricultural Revolution’: A Case Study from Stafford, England, c. AD 800– 1200. European Journal of Archaeology, 23: 585–609. https://doi.org/10.1017/eaa.2020.6

Hamilton, J. & R. Thomas. 2012. Pannage, Pulses and Pigs: Isotopic and Zooarchaeological Evidence for Changing Pig Management Practices in Later Medieval England. Medieval Archaeology, 56: 234–259. https://doi.org/10.1179/0076609712Z.0000000008

Hedges, R.E.M., R.E. Stevens & M.P. Richards. 2004. Bone as a stable isotope archive for local climatic information. Quaternary Science Reviews, 23: 959–65. https://doi.org/10.1016/j.quascirev.2003.06.022

Holmes, G. (ed.). 1988. The Oxford History of Medieval Europe. Oxford: Oxford University Press.

Hughes, S.S., A.R. Millard, C.A. Chenery, G. Nowell & D.G. Pearson. 2018. Isotopic analysis of burials from the early Anglo-Saxon cemetery at Eastbourne, Sussex, U.K. Journal of Archaeological Science: Reports, 19: 513–525. https://doi.org/10.1016/j.jasrep.2018.03.004

Jarman, C.L., M. Biddle, T. Higham & C. Bronk Ramsey. 2018. The Viking Great Army in England: new dates from the Repton charnel. Antiquity, 92: 183–199. https://doi.org/10.15184/aqy.2017.196

Lamb, A.L., J.E. Evans, R. Buckley & J. Appleby. 2014. Multi-isotope analysis demonstrates significant lifestyle changes in King Richard III. Journal of Archaeological Science, 50: 559–565. https://doi.org/10.1016/j.jas.2014.06.021

Lee-Thorp, J.A. 2008. On isotopes and old bones. Archaeometry, 50: 925–950. https://doi.org/10.1111/j.1475-4754.2008.00441.x

Leng, M.J. (ed.). 2006. Isotopes in Palaeoenvironmental Research. Dordrecht: Springer.

Lewit, T. 2009. Pigs, presses and pastoralism: farming in the fifth to sixth centuries AD. Early Medieval Europe, 17: 77–91. https://doi.org/10.1111/j.1468-0254.2009.00245.x

Lightfoot, E. & T.C. O’connell. 2016. On the Use of Biomineral Oxygen Isotope Data to Identify Human Migrants in the Archaeological Record: Intra-Sample Variation, Statistical Methods and Geographical Considerations. PLoS ONE, 11: e0153850. https://doi.org/10.1371/journal.pone.0153850

Lopez-Costas, O. & G. MüLdner. 2016. Fringes of the empire: Diet and cultural change at the Roman to post-Roman transition in NW Iberia. American Journal of Physical Anthropology, 161: 141–154. https://doi.org/10.1002/ajpa.23016

Mackinnon, M. 2019. Consistency and change: zooarchaeological investigation of Late Antique diets and husbandry techniques in the Mediterranean region. Antiquité Tardive, 27: 135–148. https://doi.org/10.1484/J.AT.5.119548

Meier-Augenstein, F. (ed.). 2010. Stable Isotope Forensics: An Introduction to the Forensic Application of Stable Isotope Analysis. Chichester: Wiley.

Montanari, M. 1988. Alimentazione e Cultura nel Medioevo. Rome: Laterza.

Pearson, K.L. 1997. Nutrition and the Early-Medieval Diet. Speculum, 72: 1–32. https://doi.org/10.2307/2865862

Reitsema, L.J., T. Kozłowski & D. Makowiecki. 2013. Humane environment interactions in medieval Poland: a perspective from the analysis of faunal stable isotope ratios. Journal of Archaeological Science, 40: 3636–3646. https://doi.org/10.1016/j.jas.2013.04.015

Richards, M.P. & K. Britton. (ed.) 2020. Archaeological Science: An Introduction. Cambridge: Cambridge University Press.

Rippon S., A. Wainwright & C. Smart. 2014. Farming Regions in Medieval England: The Archaeobotanical and Zooarchaeological Evidence. Medieval Archaeology, 58: 195–255. https://doi.org/10.1179/0076609714Z.00000000036

Roberts, P., R. Fernandes, O.E. Craig, T. Larsen, A. Lucquin, J. Swift & J. Zech. 2018. Calling all archaeologists: guidelines for terminology, methodology, data handling, and reporting when undertaking and reviewing stable isotope applications in archaeology. Rapid Communications in Mass Spectrometry, 32: 361–372. https://doi.org/10.1002/rcm.8044

Roffey, S., K. Tucker, K. Filipel-ogden, J. Montgomery, J. Cameron, T. O’connell, J. Evans, P. Marter & G.M. Taylor. 2017. Investigation of a medieval pilgrim burial excavated from the leprosarium of St Mary Magdalen Winchester, UK. PLoS Neglected Tropical Diseases 11: e0005186. https://doi.org/10.1371/journal.pntd.0005186

Salesse, K., R. Fernandes, X. De Rocheforte, J. Brůžek, D. Castex & É. Dufour. 2018. IsoArcH.eu: An open-access and collaborative isotope database for bioarchaeological samples from the Graeco-Roman world and its margins. Journal of Archaeological Science: Reports 19: 1050–1055. https://doi.org/10.1016/j.jasrep.2017.07.030

Salvadori, F. 2019. The transition from late antiquity to early Middle Ages in Italy. A zooarchaeological perspective. Quaternary International, 499: 35–48. https://doi.org/10.1016/j.quaint.2018.06.040

Styring, A.K., M. Ater, Y. Hmimsa, R., Fraser, H., Miller, R., Neef, J.A., Pearson & A. Boogard. 2016. Disentangling the effect of farming practice from aridity on crop stable isotope values: A present-day model from Morocco and its application to early farming sites in the eastern Mediterranean. The Anthropocene Review, 3: 2–22. https://doi.org/10.1177%2F2053019616630762

Vogel, J.C. & N.J. Van Der Merwe. 1977. Isotopic Evidence for Early Maize Cultivation in New York State. American Antiquity, 42: 238–242. http://www.jstor.org/stable/278984

Wang X., P. Roberts, Z. Tang, S. Yang, M. Storozum, M. Groß & R. Fernandes 2021. The Circulation of Ancient Animal Resources Across the Yellow River Basin: A Preliminary Bayesian Re-evaluation of Sr Isotope Data From the Early Neolithic to the Western Zhou Dynasty. Frontiers in Ecology and Evolution, 9: 583301. https://doi.org/10.3389/fevo.2021.583301

Wickham, C. 2016. Medieval Europe. New Haven: Yale University Press.

Wilkin, S. ET AL. 2020. Economic Diversification supported the growth of Mongolia’s Nomadic empires. Scientific reports, 10: 3916. https://doi.org/10.1038/s41598-020-60194-0

Winklerová, D. 2011. Zooarchaeological and archaeobotanical indicators for aspects of diet in medieval Kingdom of Bohemia, in J. Klápště & P. Sommer (ed.) Processing, storage, distribution of food: food in the medieval rural environment: 421–429. Turnhout: Brepols.

Witcher, R. 2016. Agricultural Production in Roman Italy, in A. Cooley (ed.), A Companion to Roman Italy: 459–482. Chichester: Wiley-Blackwell.

