## Supplementary figures and images for "Presenting the *Compendium Isotoporum Medii Aevi* (CIMA) and Bayesian Case Studies"

### Supp. Info 3

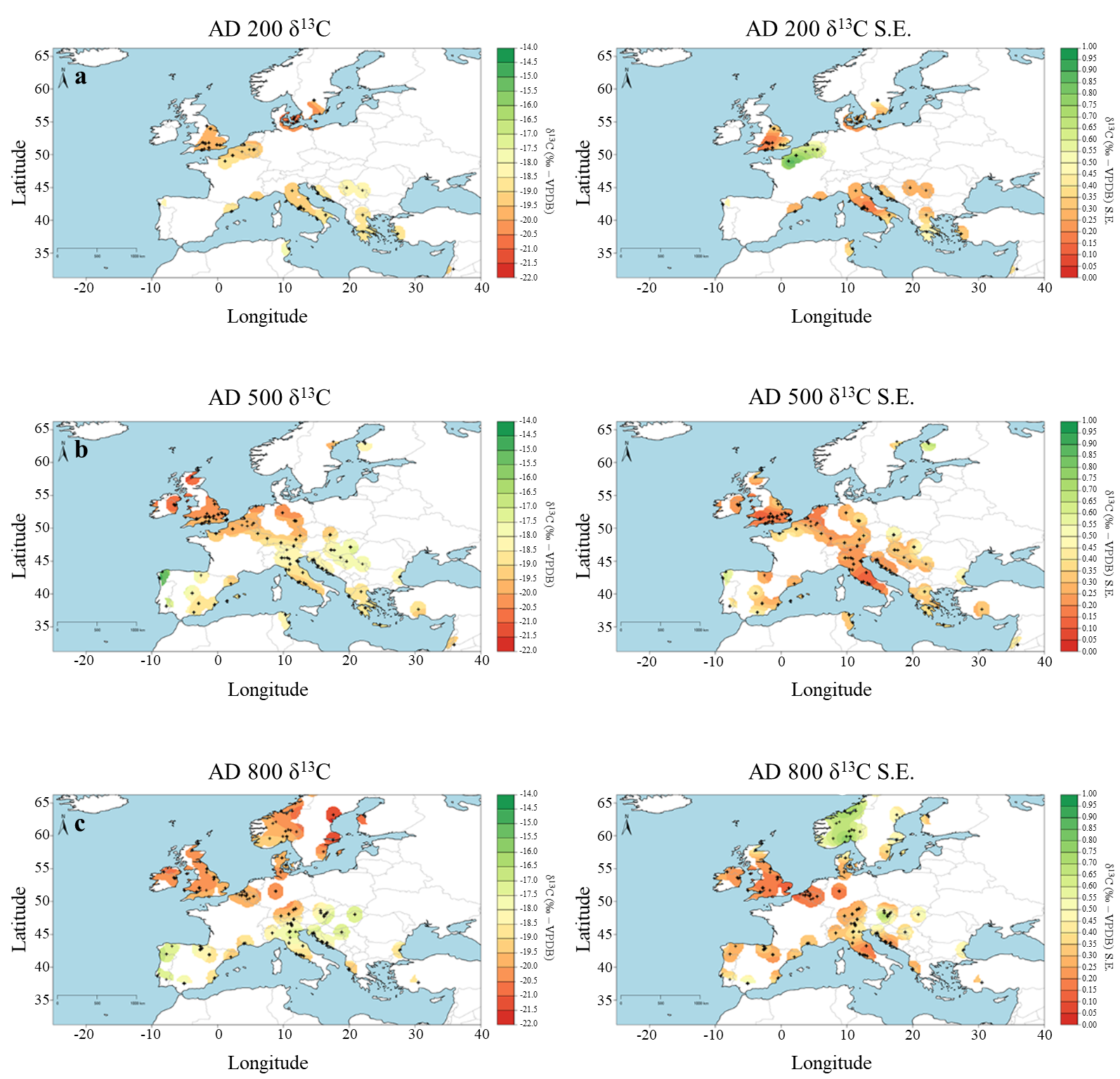

### Supp. Info 4

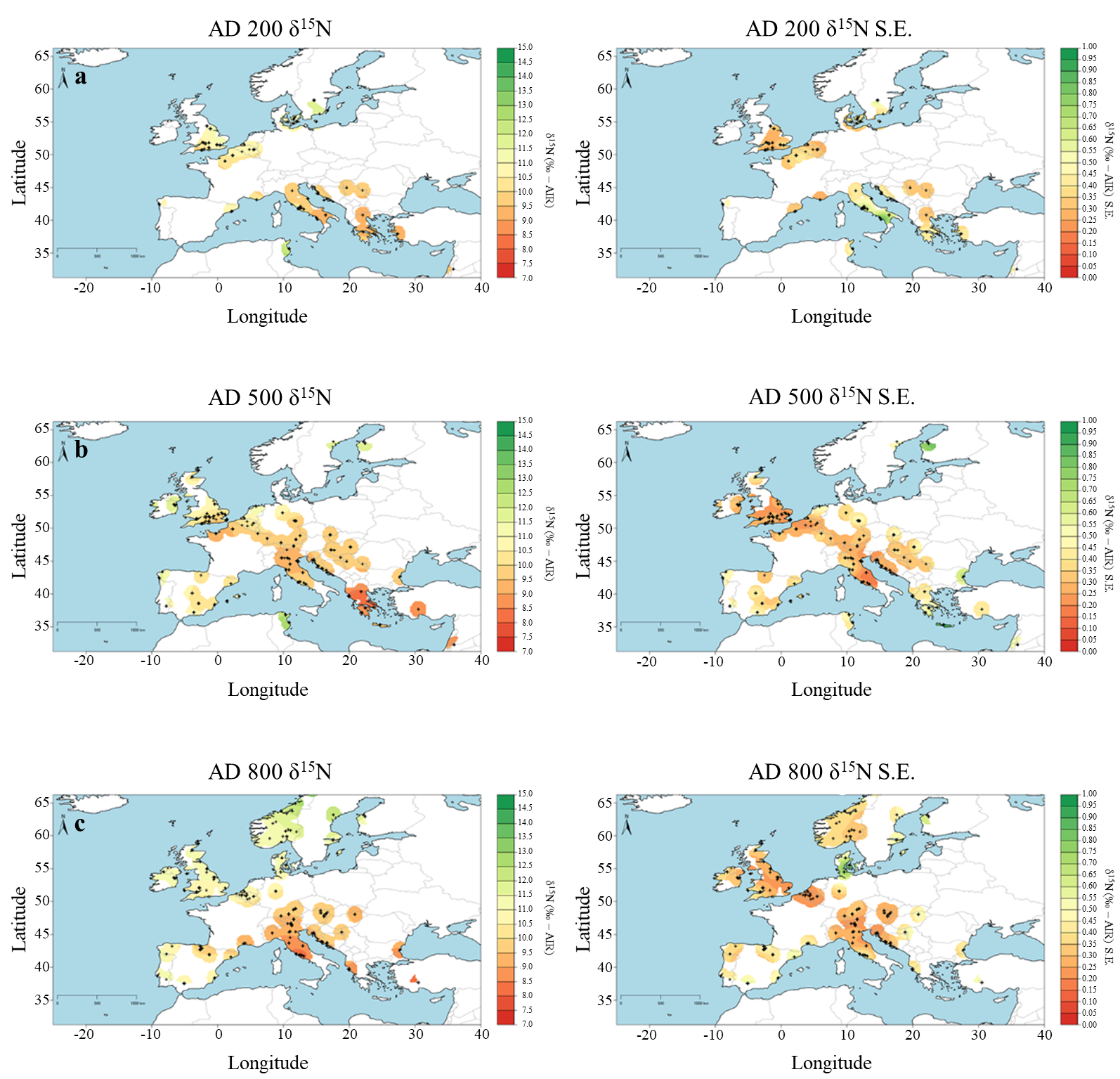
