## Supplementary material for "Presenting the *Compendium Isotoporum Medii Aevi* (CIMA) and Bayesian Case Studies": Supp. Info 2

### Supplementary Information File S2: Bayesian modelling

#### Pandora & IsoMemo modelling

The Bayesian case studies described in the main text employed modelling tools (TimeR, AverageR, OperatoR, KernelTimeR and LocateR) developed within the Pandora & IsoMemo initiatives (https://pandoraapp.earth/). Below we give a description of these models.

*TimeR*

'TimeR' is a Bayesian geostatistical model that estimates the expected value of a "dependent" variable across time and space. The underlying model formula is:

Y_i = g(long_i, lat_i, time_i) + gamma_site_i + epsilon_i

gamma_site_i is a random effect with ~ N(0, tau^2)

where g()- is a three dimensional (spatial coordinates plus time) smooth function using a so-called thin plate regression spline ("tprs", Wood 2003) and epsilon follows a normal distribution N(0, sigma^2). As the time variable is measured with uncertainty, a Metropolis-Hastings step is employed to account for it (further details given in Groß (2016)). The model was employed in case-study 2 to model spatio-temporal variations in human bone collagen isotopic values so that shifts in human diets for early medieval Europe and for the area centred Rome could be assessed (Fig. 7; also S3-S4). Filtered data for analyses consisted of non-elite adult individuals with bone and tooth collagen C:N atomic ratios within the acceptance interval given by van Klinken (1999).

*AverageR*

'AverageR' is a Bayesian geostatistical model that estimates the expected value of the "dependent" variable across space. 'AverageR' is equivalent to 'TimeR' but does not include time dependence (Cubas *et al.* 2020). The model was employed in case-study 1 to map the spatial distribution of $\delta^{13}C$ and $\delta^{15}N$ mean and prediction error values (double the square root of the following sum: the square of the predicted standard error of the mean plus the square of the predicted population standard deviation) for medieval domesticated animals (Fig. 4) and in case-study 3, to establish reference baselines for mobility studies (see below, LocateR). For the animals, we modelled separately isotopic values for cattle and sheep/goat *versus* pig and chicken with acceptable C/N atomic ratios van Klinken (1999).

##### *OperatoR*

OperatoR was used to plot the differences among maps generated using AverageR. This difference was calculated by subtracting two-dimensional posterior estimates of the time sliced g()-smooth, i.e. the expected values for all considered locations for a fixed date. To establish the significance of the difference the standard error of the difference was computed and tested against a null hypothesis of no difference. OperatoR was used in case-study 2 to show the temporal differences in early medieval diets for two different time-slices (Fig. 5-6). 'TimeR' was used to generate spatial models in three different time-slices (AD 200-500-800) followed by the use of OperatoR to map differences.

##### *KernelTimeR*

'KernelTimeR' is a 3-dimensional spatiotemporal kernel density estimator (e.g. Wand & Jones 1994). It was employed in case-study 3 to show research gaps in early medieval isotopic studies and to assess temporal variability in human mobility patterns (Fig. 8). Mobile *versus* non-mobile individuals were identified by comparing individual values with a $^{87}Sr/^{86}Sr$ and $\delta^{18}O_{phosphate}$ tooth enamel baseline for Britain (Evans *et al.* 2010; Pellegrini *et al.* 2016) modelled using 'AverageR'. Individuals were classified as mobile if their $^{87}Sr/^{86}Sr$ or $\delta^{18}O$ tooth values were outside the 2-sigma range for local baselines.

##### *LocateR*

Given a 2-dimensional grid of locations and corresponding estimates of mean and residual error from the 'TimeR' or 'AverageR' models, for a new given value of the dependent variable, 'LocateR' assigns a density value to each location (Wang *et al.* 2021). The model was employed in case-study 3 to show probable dwelling locations (and likely places of origin for early formed teeth) for four different medieval individuals buried in England. For this example, we used'AverageR' to establish a baseline for water $\delta^{18}O$ using the Cluster-based Water Isotope Prediction Model (RCWIP), which relies on modern measurements from the Global Network of Isotopes in Precipitation (GNIP) (Terzer *et al.* 2013) and a $^{87}Sr/^{86}Sr$ and $\delta^{18}O_{phosphate}$ tooth enamel baseline for Britain (Evans *et al.* 2010; Pellegrini *et al.* 2016).

#### ReSources

ReSources is a software used to define Bayesian mixing models that can be employed for isotope-based diet reconstruction. This software is an upgraded version of the Bayesian software FRUITS (Fernandes *et al.* 2014). We employed the model to quantify caloric and macronutrient contribution of each food source within human diets in the area of Rome in three time-slices: AD 200, AD 500 and AD 800. The model employs a random effects structure on the categorical covariate "Time" with levels AD 200, AD 500 and AD 800. In contrast to a model without covariates, the dirichlet prior values of the source contribution parameters ("alpha") are not fixed but come from a distribution with common mean and standard deviation for each factor level (AD 200, 500, 800) and source group.

Although the model has been employed in recent years to reconstruct past human diets (e.g. Fernandes *et al.* 2015; Bownes *et al.* 2017; Varano *et al.* 2020), the chosen food categories were often very broad (e.g. following chemical and/or habitat classifications such as $C_3/C_4$ plants, $C_3$ animals, marine animals, etc.). In our case-study, we relied on archaeobotanical, zooarchaeological and historical evidence to define the following main food group for Roman and medieval populations at Romes: Wheat; Barley; $C_4$ Cereals (i.e. millet or sorghum); Pulses; Other $C_3$ Plants (i.e. Vegetables, fruit, nuts); Cattle; Sheep/Goat; Pig; Poultry; Marine Sources; Freshwater Sources. Missing from the above, given a lack of isotopic references, is olive oil whose contributions will likely fall among the $C_3$ plants.

Human mean isotopic values for Rome (Lat. 41.9; Long. 12.5) were modelled using TimeR from bone collagen measurements available at CIMA and IsoArcH (Salesse *et al.* 2018) databases. We considered three separate time slices (AD 200, 500, and 800) with the following results: AD 200: $\delta^{13}C$=-19.14±0.11‰, $\delta^{15}N$=10.08±0.11‰; AD 500: $\delta^{13}C$=-19.24±0.2‰, $\delta^{15}N$=9.57±0.23‰; AD 800: $\delta^{13}C$=-19.05±0.38‰, $\delta^{15}N$=8.6±0.23‰. As for the isotopic food baseline (tab. S2.1 below), wheat, barley and pulses isotopic values were obtained from the O'Connell *et al.* (2019) publication on Portus. Given the lack of direct isotopic measurements on archaeological fruit and vegetables, isotopic values for the 'other $C_3$ plants' were calculated by applying a trophic isotopic offset correction to herbivore mean bone collagen carbon and nitrogen stable isotope values ($\delta^{13}C$=-4‰;

$\delta^{15}N$=-3.5, Lee-Thorp *et al.* 1989; Hedges & Reynard 2007) between these and $C_3$ plants. Lacking coeval isotopic values on $C_4$ cereals, we employed isotopic measurements from two sites in Bronze Age Greece (Nitsch *et al.* 2017), as reported in the IsoArch database. As for food products from cattle, sheep/goat and pig the isotopic references given below were modelled using 'TimeR' for Rome (Lat. 41.9; Long. 12.5). For poultry, given a limited amount of data, we calculated the mean and standard deviation for isotopic values available from CIMA (Baldoni *et al.* 2016; Buonincontri *et al.* 2017; Maxwell 2019; O'Connell *et al.* 2019; Rolandsen *et al.* 2020; Gismondi *et al.* 2020; Paladin *et al.* 2020; Riccomi *et al.* 2020; Varano *et al.* 2020; Viva *et al.* 2021). A similar approach was employed for Mediteranean marine sources by combining CIMA and IsoArcH data (Alexander *et al.* 2015; Sandias & Müldner 2015; O'Connell *et al.* 2019; Gismondi *et al.* 2020; Ma *et al.* 2021). For freshwater sources, we collected medieval measurements from Italy (Riccomi *et al.* 2020), Switzerland (Häberle *et al.* 2016) and France (Mion *et al.* 2019). Food sources isotopic values were the following:

| | AD 200 $\delta^{13}C$ | AD 500 $\delta^{13}C$ | AD 800 $\delta^{13}C$ |
|---|---|---|---|
| Wheat | -23.18±0.87‰ | -22.07±0.96‰ | -22.45±0.35‰ |
| Barley | -24.4±0.7‰ | -22±0.4‰ | -23.46±0.15‰ |
| $C_4$ Cereals | -10.4±0.27‰ | -10.4±0.27‰ | -10.4±0.27‰ |
| Pulses | -24±1.3‰ | -26.5±0.4‰ | -21.87±0.75‰ |
| Other $C_3$ Plants | -24.3±0.4‰ | -24±0.3‰ | -24±0.3‰ |
| Cattle | -20.2±0.34‰ | -19.76±0.15‰ | -19.75±0.29‰ |
| Sheep/Goat | -20.35±0.19‰ | -20.31±0.23‰ | -20.25±0.17‰ |
| Pig | -20.15±0.12‰ | -19.88±0.07‰ | -20.16±0.11‰ |
| Poultry | -18.7±2‰ | -18.7±2‰ | -18.7±2‰ |
| Marine Sources | -11.3±3‰ | -11.3±3‰ | -11.3±3‰ |
| Freshwater Sources | -23.39±1.3‰ | -23.39±1.3‰ | -23.39±1.3‰ |

| | AD 200 $\delta^{15}N$ | AD 500 $\delta^{15}N$ | AD 800 $\delta^{15}N$ |
|---|---|---|---|
| Wheat | 8.53±3.11‰ | 6.76±3.84‰ | 4.3±0.28‰ |
| Barley | 6.1±1.3‰ | 6.9±0.4‰ | 5.63±0.97‰ |

| | | | |
|---|---|---|---|
| $C_4$ Cereals | 6.8±2.74‰ | 6.8±2.74‰ | 6.8±2.74‰ |
| Pulses | 2.5±0.6‰ | 2.5±0.6‰ | 2.4±0.46‰ |
| Other $C_3$ Plants | 2.1±0.6‰ | 2±0.8‰ | 2.3±0.6‰ |
| Cattle | 5.96±0.41‰ | 5.99±0.77‰ | 6.27±0.43‰ |
| Sheep/Goat | 5.29±0.41‰ | 4.94±0.25‰ | 5.32±0.39‰ |
| Pig | 6.34±0.24‰ | 6.13±0.16‰ | 5.98±0.34‰ |
| Poultry | 7.6±2.1‰ | 7.6±2.1‰ | 7.6±2.1‰ |
| Marine Sources | 10.7±1.9‰ | 10.7±1.9‰ | 10.7±1.9‰ |
| Freshwater Sources | 7.98±1.88‰ | 7.98±1.88‰ | 7.98±1.88‰ |

Tab. S2.1. Isotopic values for each food source and time-slice employed in Bayesian dietary modelling.

Within the Bayesian mixing model we employed macronutrient (carbs/lipids vs. protein) composition values as reported in Fernandes *et al.* 2015 but with doubled uncertainty values. These were for plant foods: protein: 10±5%; carbs/lipids 90±5%%), terrestrial animals: protein: 30±5%; carbs/lipids 70±5%, and aquatic animals: protein: 65±10%; carbs/lipids 35±10%. To obtain $\delta^{13}C$ and $\delta^{15}N$ values of protein and carbohydrates/lipids food components we applied offset corrections between the measured material and the edible nutritional component of the respective food group (Tab. S2.2, below). We applied the following offset corrections with uncertainties for macronutrient isotopic values rounded up to multiples of 0.5 (Fernandes *et al.* 2015; updated in Soncin *et al.* 2021): Plants: $\Delta^{13}C_{protein\text{-}bulk}$=-2‰, $\Delta^{13}C_{carbohydrates\text{-}bulk}$=+0.5‰, $\Delta^{15}N_{protein\text{-}bulk}$=0‰; terrestrial animals: $\Delta^{13}C_{protein\text{-}collagen}$= -2‰, $\Delta^{13}C_{lipids\text{-}collagen}$= -8‰, $\Delta^{15}N_{protein\text{-}collagen}$=0‰; aquatic animals: $\Delta^{13}C_{protein\text{-}collagen}$=-1‰, $\Delta^{13}C_{lipids\text{-}collagen}$=-7‰, $\Delta^{15}N_{protein\text{-}collagen}$=+1.5‰). Below a list of isotopic reference values employed in modelling:

| | AD 200 $\delta^{13}C$ Protein | AD 200 $\delta^{13}C$ Lipids/Carbohydrates | AD 200 $\delta^{15}N$ Protein |
|---|---|---|---|
| Wheat | -25.2±2‰ | -22.7±2‰ | 8.5±4.5‰ |
| Barley | -26.4±2‰ | -23.9±2‰ | 6.1±2.5‰ |

| | | | |
|---|---|---|---|
| $C_4$ Cereals | -12.4±1.5‰ | -9.9±1.5‰ | 6.8±4.0‰ |
| Pulses | -26.0±2.5‰ | -23.5±2.5‰ | 2.5±2‰ |
| Other $C_3$ Plants | -26.3±1.5‰ | -23.8±1.5‰ | 2.1±2‰ |
| Cattle | -22.2±1.5‰ | -28.2±1.5‰ | 6±1.5‰ |
| Sheep/Goat | -22.4±1.5‰ | -28.4±1.5‰ | 5.3±1.5‰ |
| Pig | -22.2±1.5‰ | -28.2±1.5‰ | 6.3±1.5‰ |
| Poultry | -20.7±3‰ | -26.7±3‰ | 7.6±3.5‰ |
| Marine Sources | -12.3±4‰ | -18.3±4‰ | 12.2±3‰ |
| Freshwater Sources | -24.4±2.5‰ | -30.4±2.5‰ | 9.5±3‰ |

| | AD 500 $\delta^{13}C$ Protein | AD 500 $\delta^{13}C$ Lipids/Carbohydrates | AD 500 $\delta^{15}N$ Protein |
|---|---|---|---|
| Wheat | -24.1±2‰ | -21.6±2‰ | 6.8±5‰ |
| Barley | -24.0±1‰ | -21.5±1‰ | 6.9±1‰ |
| $C_4$ Cereals | -12.4±1.5‰ | -9.9±1.5‰ | 6.8±4.0‰ |
| Pulses | -28.5±1‰ | -26±1‰ | 2.5±2‰ |
| Other $C_3$ Plants | -26.0±1.5‰ | -23.5±1.5‰ | 2±2‰ |
| Cattle | -21.8±1.5‰ | -27.8±1.5‰ | 6.0±2‰ |
| Sheep/Goat | -22.3±1.5‰ | -28.3±1.5‰ | 4.9±1.5‰ |
| Pig | -21.9±1.5‰ | -27.9±1.5‰ | 6.1±1.5‰ |
| Poultry | -20.7±3‰ | -26.7±3‰ | 7.6±3.5‰ |
| Marine Sources | -12.3±4‰ | -18.3±4‰ | 12.2±3‰ |
| Freshwater Sources | -24.4±2.5‰ | -30.4±2.5‰ | 9.5±3‰ |

| | AD 800 $\delta^{13}C$ Protein | AD 800 $\delta^{13}C$ Lipids/Carbohydrates | AD 800 $\delta^{15}N$ Protein |
|---|---|---|---|
| Wheat | -24.5±1.5‰ | -22.0±1.5‰ | 4.3±1.5‰ |
| Barley | -25.5±1.5‰ | -23.0±1.5‰ | 5.6±2‰ |

| | | | |
|---|---|---|---|
| $C_4$ Cereals | -12.4±1.5‰ | -9.9±1.5‰ | 6.8±4.0‰ |
| Pulses | -23.9±2‰ | -21.4±2‰ | 2.4±1.5‰ |
| Other $C_3$ Plants | -26±1.5‰ | -23.5.0±1.5‰ | 2.3±2‰ |
| Cattle | -21.8±1.5‰ | -27.8±1.5‰ | 6.3±1.5‰ |
| Sheep/Goat | -22.3±1.5‰ | -27.8±1.5‰ | 5.3±1.5‰ |
| Pig | -22.2±1.5‰ | -28.2±1.5‰ | 6±1.5‰ |
| Poultry | -20.7±3‰ | -26.7±3‰ | 7.6±3.5‰ |
| Marine Sources | -12.3±4‰ | -18.3±4‰ | 12.2±3‰ |
| Freshwater Sources | -24.4±2.5‰ | -30.4±2.5‰ | 9.5±3‰ |

Tab.S2.2. Corrected isotopic values as employed in the model according to each food group, time-slice and dietary route.

The Bayesian mixing model also accounted for isotopic offsets between diet and human tissues and dietary routing mechanisms. Bone collagen $\delta^{15}N$ was assumed to fully derive from dietary protein with an isotopic offset of 5.5±0.5‰ (Fernandes *et al.* 2015). For bone collagen $\delta^{13}C$ we considered an offset of 4.8±0.5‰ and that the isotopic signal was routed by 74±4% from dietary protein and 26±4% from carbohydrates/lipids (Fernandes *et al.* 2012).

To improve the precision of dietary estimates we employed prior dietary information from zooarchaeological, archaeobotanical and historical evidence relative to ancient Rome (Fig. 8). We considered that, in farming societies plants were the main caloric sources for lower status individuals. We set the following constraint in the Bayesian model: Wheat+barley+$C_4$ cereals+pulses>57% of the caloric contribution. The value is the mean caloric contribution from starches consumed in Mediterranean countries between 1960–1965 (table 14.2 reported in Garnsey 1998: 231).

We collected archaeobotanical evidence from central and southern Italy (Sadori & Susanna 2005; Caracuta & Fiorentino 2009; Van der Noort *et al.* 2009; Murphy *et al.* 2013; Robinson & Rowan 2015; Buonincontri *et al.* 2017), and from this ordered plant starch dietary contributions as follows: Wheat>Pulses>Barley>Millet. This data plus historical sources (Montanari 1988) also revealed a temporal decrease in wheat. The consumption of $C_4$ cereals increased during the medieval period for rural Italian populations (Montanari 1988). During the Late Roman and early medieval period, the consumption of animal food sources decreased and populations relied increasingly on cereals, legumes, and other plants (Montanari 1988).

The zooarchaeological evidence (King 1999; Minniti 2005; MacKinnon 2019) shows that pig remains were the most abundant for all periods although this decreased through time in favour of sheep/goat and, to a lesser extent, cattle.

Dietary modelling results are shown in Fig. 8 and summarised in Supplementary Information file S5. The latter also includes dietary estimates for modern Italians. For this we employed data from FAO food balance sheets (http://www.fao.org/faostat/en/#data/FBS) for Italy during 2018 and from 1994–96 (Turrini *et al.* 1999). A comparison was made only for food sources significantly available in Roman and early medieval Italy. However, we also list non-filtered food estimates for modern Italians.

#### Supplementary Information File S2 References


Alexander, M.M., C.M. Gerrard, A. Gutiérrez & A.R. Millard. 2015. Diet, Society, and Economy in Late Medieval Spain: Stable Isotope Evidence from Muslims and Christians From Gandía, Valencia. *American Journal of Physical Anthropology*, 156: 263–273. https://doi.org/10.1002/ajpa.22647

Baldoni, M. et al. 2016. Archaeo-biological reconstruction of the Italian medieval population of Colonna (8th–10th centuries CE). *Journal of Archaeological Science: Reports*, 10: 483–494. https://doi.org/10.1016/j.jasrep.2016.11.013

Bocherens, H., M. Fizet, A., Mariotti, C., Olive, G., Bellon & D., Billiou. 1991. Application de la biogeochimie isotopique (13C, 15N) a la déetermination du régime alimentaire des populations humains et animales durante les périodes antique et médiévale. *Archives des Sciences – Université de Genève*: 44: 329–340.

Bownes, J.M., P.L. Ascough, G.T. Cook, I. Murray & C. Bonsall. 2017. Using Stable Isotopes and a Bayesian Mixing Model (FRUITS) to Investigate Diet at the Early Neolithic Site of Carding Mill Bay, Scotland. *Radiocarbon*, 59: 1275–1294. https://doi.org/10.1017/RDC.2017.39

BUONINCONTRI M.P., A. PECCI, G. DI PASQUALE, P. RICCI & C. LUBRITTO. 2017. Multiproxy approach to the study of Medieval food habits in Tuscany (central Italy). *Archaeological and Anthropological Sciences*, 9: 653–671. https://doi.org/10.1007/s12520-016-0428-7

CARACUTA, V. & G. FIORENTINO. 2009. L'analisi archeobotanica nell'insediamento di Faragola (FG): il paesaggio vegetale tra spinte antropiche e caratteristiche ambientali, tra tardoantico e altomedioevo, in G. Volpe & P. Favia (ed.) *Atti del V Congresso Nazionale di Archeologia Medievale*. 371–377. Firenze: All'Insegna del Giglio.

CUBAS, M. ET AL. 2020. Latitudinal gradient in dairy production with the introduction of farming in Atlantic Europe. *Nature communications*, 11: 2036. https://doi.org/10.1038/s41467-020-15907-4

EVANS, J., J. MONTGOMERY, G. WILDMAN & N. BOULTON. 2010. Spatial variations in biosphere $^{87}Sr/^{86}Sr$ in Britain. *Journal of the Geological Society*, 167: 1–4. https://doi.org/10.1144/0016-76492009-090

FERNANDES, R., M.J. NADEAU & P.M. GROOTES. 2012. Macronutrient-based model for dietary carbon routing in bone collagen and bioapatite. *Archaeological and Anthropological Sciences*, 4: 291–301. https://doi.org/10.1007/s12520-012-0102-7

FERNANDES, R, A.R. MILLARD, M. BRABEC, M.J. NADEAU & P. GROOTES. 2014. Food reconstruction using isotopic transferred signals (FRUITS): a Bayesian model for diet reconstruction. *PloS ONE*, 9: e87436. https://doi.org/10.1371/journal.pone.0087436

FERNANDES, R, P. GROOTES, M.J. NADEAU & O. NEHLICH. 2015 Quantitative diet reconstruction of a Neolithic population using a Bayesian mixing model (FRUITS): the case study of Ostorf (Germany). *American Journal of Physical Anthropology*, 158: 325–340. https://doi.org/10.1002/ajpa.22788

GARNSEY, P. (1998). *Cities, Peasants and Food in Classical Antiquity. Essays in Social and Economic History.* Cambridge: Cambridge University Press.

GISMONDI, A. ET AL. 2020. A multidisciplinary approach for investigating dietary and medicinal habits of the Medieval population of Santa Severa (7th-15th centuries, Rome, Italy). *PLoS One*, 15: e0227433. https://doi.org/10.1371/journal.pone.0227433

GROß, M. 2016. Modeling body height in prehistory using a spatio-temporal Bayesian errors-in variables model. *AStA Advances in Statistical Analysis*, 100: 289–311. https://doi.org/10.1007/s10182-015-0260-x

HÄBERLE, S., B.T. FULLER, O. NEHLICH, W. VAN NEER, J. SCHIBLER, J. & H. HÜSTER PLOGMANN. 2016. Inter- and intraspecies variability in stable isotope ratio values of archaeological freshwater fish remains from Switzerland (11th–19th centuries AD). *Environmental Archaeology*, 21: 119–132. https://doi.org/10.1179/1749631414Y.0000000042

HEDGES, R.E.M. & L.M. REYNARD. 2007. Nitrogen isotopes and the trophic level of humans in archaeology. *Journal of Archaeological Science*, 34: 1240–1251. https://doi.org/10.1016/j.jas.2006.10.015

KING, A. 1999. Diet in the Roman world: a regional intersite comparison. *Journal of Roman Archaeology*, 12: 168–202.

LEE-THORP, J.A., J.C. SEALY & N.J. VAN DER MERWE. 1989. Stable carbon isotope ratio differences between bone collagen and bone apatite, and their relationship to diet. *Journal of archaeological science*, 16: 585–599. https://doi.org/10.1016/0305-4403(89)90024-1

MA, Y., R. BOCKMANN, S.T. STEVENS, S. ROUDESLI-CHEBBI, A. AMARO, A. BROZOU, B.T. FULLER & M.A. MANNINO. 2021. Isotopic reconstruction of diet at the Vandalic period (c. 5th – 6th centuries AD) Theodosian Wall cemetery at Carthage, Tunisia. *International Journal of Osteoarchaeology*, 31: 393–405. https://doi.org/10.1002/oa.2958

MACKINNON, M. 2019. Consistency and change: zooarchaeological investigation of Late Antique diets and husbandry techniques in the Mediterranean region. *Antiquité Tardive*, 27: 135–148. https://doi.org/10.1484/J.AT.5.119548

MAXWELL, A. 2019. Exploring Variations in Diet and Migration from Late Antiquity to the Early Medieval Period in the Veneto, Italy: A Biochemical Analysis. Unpublished PhD dissertation: University of South Florida.

MINNITI, C. 2005. L'approvvigionamento alimentare a Roma nel Medioevo: analisi dei resti faunistici provenienti dalle aree di scavo della Crypta Balbi e di Santa Cecilia, in I. Fiore, G. Malerba & S. Chilardi (ed.) *Atti del III Convegno Nazionale di Archeozoologia*: 469–492. Roma: Istituto poligrafico e Zecca dello Stato.

MION, L., E. HERRSCHER, G. ANDRÉ, J. HERNANDEZ, R. DONAT, M, FABRE, V. FOREST & D.C. SALAZAR-GARCÍA. 2019. The influence of religious identity and socio-economic status on diet over time, an example from medieval France. *Archaeological and Anthropological Sciences*, 11: 3309–3327. https://doi.org/10.1007/s12520-018-0754-z

MONTANARI, M. 1988. *Alimentazione e Cultura nel Medioevo*. Rome: Laterza.

MURPHY, C., G. THOMPSON & D.Q. FULLER. 2012. Roman food refuse: urban archaeobotany in Pompeii, Regio VI, Insula 1. *Vegetation History and Archaeobotany*, 22: 409–419. https://doi.org/10.1007/s00334-012-0385-8

NITSCH, E. ET AL. 2017. A bottom-up view of food surplus: using stable carbon and nitrogen isotope analysis to investigate agricultural strategies and diet at Bronze Age Archontiko and Thessaloniki Toumba, northern Greece. *World Archaeology*, 49: 105–137. https://doi.org/10.1080/00438243.2016.1271745

O'CONNELL, T.C., R.M. BALLANTYNE, S. HAMILTON-DYER, E. MARGARITIS, S. OXFORD, W. PANTANO, M. MILLETT & S.J. KEAY. 2019. Living and dying at the *Portus Romae*. *Antiquity*, 93: 719–734. https://doi.org/10.15184/aqy.2019.64

PALADIN, A., N. MOGHADDAM, A.E. STAWINOGA, I., SIEBKE, V. DEPELLEGRIN, U. TECCHIATI, S. LÖSCH & A. ZINK. 2020. Early medieval Italian Alps: reconstructing diet and mobility in the valleys. *Archaeological and Anthropological Science*, 12: 82. https://doi.org/10.1007/s12520-019-00982-6

PELLEGRINI, M., J. POUNCETT, M. JAY, M.P. PEARSON & M.P. RICHARDS. Tooth enamel oxygen "isoscapes" show a high degree of human mobility in prehistoric Britain. *Scientific Reports*, 6: 34986. https://doi.org/10.1038/srep34986

RICCOMI, G., S. MINOZZI, J. ZECH, F. CANTINI, V. GIUFFRA & P. ROBERTS, P. 2020. Stable isotopic reconstruction of dietary changes across Late Antiquity and the Middle Ages in Tuscany. *Journal of Archaeological Science: Reports* 33: 102546. https://doi.org/10.1016/j.jasrep.2020.102546

ROBINSON, M. & E. ROWAN. 2015. Roman Food Remains in Archaeology and the Contents of a Roman Sewer at Herculaneum, in J. Wilkins & R. Nadeau (ed.) *A Companion to Food in the Ancient World*: 105–115. Chichester: Wiley-Blackwell.

ROLANDSEN, G.L., P. ARTHUR, & M. ALEXANDER. 2019. A tale of two villages: Isotopic insight into diet, economy, cultural diversity and agrarian communities in medieval (11th-15th century CE) Apulia, Southern Italy. *Journal of Archaeological Science: Reports* 28: 102009. https://doi.org/10.1016/j.jasrep.2019.102009

SADORI, L. & F. SUSANNA. 2005. Hints of economic change during the late Roman Empire period in central Italy: a study of charred plant remains from "La Fontanaccia", near Rome.

*Vegetation History and Archaeobotany*, 14: 386–393. https://doi.org/10.1007/s00334-005-0010-1

SALESSE, K., R. FERNANDES, X. DE ROCHEFORTE, J. BRŮŽEK, D. CASTEX & É. DUFOUR. 2018. IsoArcH.eu: An open-access and collaborative isotope database for bioarchaeological samples from the Graeco-Roman world and its margins. *Journal of Archaeological Science: Reports*, 19: 1050–1055. https://doi.org/10.1016/j.jasrep.2017.07.030

SANDIAS, M. & G. MÜLDNER. 2015. Diet and herding strategies in a changing environment: Stable isotope analysis of Bronze Age and Late Antique skeletal remains from Ya'amūn, Jordan. *Journal of Archaeological Science*, 63: 24–32. https://doi.org/10.1016/j.jas.2015.07.009

SONCIN, S. ET AL. 2021. High-resolution dietary reconstruction of victims of the AD 79 Vesuvius eruption at Herculaneum by compound specific isotope analysis. *Science Advances*, in press.

TERZER, S., L.I. WASSENAAR, L.J. ARAGUÁS-ARAGUÁS & P.K. AGGARWAL. 2013. Global isoscapes for $\delta^{18}O$ and $\delta^{2}H$ in precipitation: improved prediction using regionalized climatic regression models. *Hydrology and Earth System Sciences*, 17: 4713–4728. https://doi.org/10.5194/hess-17-4713-2013.

TURRINI, A., C. LECLERCQ & A. D'AMICIS. 1999. Patterns of food and nutrient intakes in Italy and their application to the development of food-based dietary guidelines. *British Journal of Nutrition*, 81: S83–S89. https://doi.org/10.1017/S0007114599001762

WAND, M. & M. JONES.1994. Multivariate plug-in bandwidth selection. *Computational Statistics*, 9: 97–116.

WANG X., P. ROBERTS, Z. TANG, S. YANG, M. STOROZUM, M. GROß & R. FERNANDES 2021. The Circulation of Ancient Animal Resources Across the Yellow River Basin: A Preliminary Bayesian Re-evaluation of Sr Isotope Data From the Early Neolithic to the Western Zhou Dynasty. *Frontiers in Ecology and Evolution*, 9: 583301. https://doi.org/10.3389/fevo.2021.583301

WOOD, S.N. 2003. Thin plate regression splines. *Journal of the Royal Statistical Society: Series B*, 65: 95–114. https://doi.org/10.1111/1467-9868.00374

VAN DER NOORT, R. ET AL.2009. Excavations at Le Mura di Santo Stefano, Anguillara Sabazia. *Papers of the British School at Rome*, 77: 159–223. https://doi.org/10.1017/S0068246200000076

VAN KLINKEN, G.J. 1999. Bone Collagen Quality Indicators for Palaeodietary and Radiocarbon Measurements. *Journal of Archaeological Science*, 26: 687–695. https://doi.org/10.1006/jasc.1998.0385

VARANO, S. ET AL. 2020. The edge of the Empire: diet characterization of medieval Rome through stable isotope analysis. *Archaeological and Anthropological Science*, 12: 196. https://doi.org/10.1007/s12520-020-01158-3

VIVA, S., P.F. FABBRI, P. RICCI, G. BIANCHI, R. HODGES & C. LUBRITTO. 2021. Project nEU-Med. The Contribution of Isotopic Analysis in the Differential Diagnosis of Anemia, the Case of the Medieval Cemetery of Vetricella (Scarlino, GR) in Tuscany. *Environmental Archaeology*. https://doi.org/10.1080/14614103.2020.1867290
